# Leg blood flow and muscle oxygenation are reduced when exercising with loads ≥40% 1RM compared to loads ≤30% 1RM due to increases in contraction induced intramuscular pressure

**DOI:** 10.64898/2026.10.24.761369

**Authors:** Matthew D. Fliss, Kyla Coates, Sam J. Hillen, Gabby Yarema, Lucas Wiens, Jem I. Arnold, Aidan Comeau, Viviana J. Shiffman, Jamie Burr, Martin MacInnis, Robert Boushel, Michael Koehle, Cameron Mitchell

**Affiliations:** The University of British Columbia; The University of British Columbia Okanagan; University of Guelph; University of Calgary; University Of British Columbia

## Abstract

The critical occluding tension (COT) is the load at which contraction-induced increases in intramuscular pressure (IMP) collapse intramuscular vasculature and impair oxygen delivery. For knee extension resistance exercise, the COT occurs at ∼30-40% 1RM, yet few studies have measured IMP alongside convective oxygen delivery. This study examined the interaction between IMP, leg blood flow, and muscle oxygenation during resistance exercise. Twelve participants completed knee extension bouts across a range of loads while vastus lateralis IMP, femoral blood flow (Doppler), blood pressure, muscle oxygenation (NIRS), and electromyography (EMG) were recorded. Mean IMP increased alongside relative load (20% 1RM=19.1(9.4) mmHg; 30% 1RM=22.4(10.4) mmHg; 40% 1RM=27.0(8.7) mmHg; 50% 1RM=31.5(11.5) mmHg; 60% 1RM=41.14(13.8) mmHg; p<0.05). Leg blood flow scaled to rest was similar between 20% and 30% 1RM (p=0.13), 40%, 50%, and 60% 1RM (p>0.05), but different between the 20-30% 1RM and 40-60% 1RM groupings (p<0.05) (20% 1RM=2.6(1.4) times rest; 30% 1RM=1.8(0.5) fold of rest; 40% 1RM=1.4(0.6) fold of rest; 50% 1RM=1.1(0.4) fold of rest; 60% 1RM=1.1(0.5) fold of rest). Muscle oxygenation was lower at 40%, 50%, and 60% 1RM compared to 20% and 30% 1RM (p<0.05). Thus, at ≤30% 1RM exercising blood flow increased well above rest, indicating preserved microvascular perfusion, whereas at ≥40% 1RM the rise in IMP was sufficient to impair perfusion and limit blood flow increases. Exercise at or above the COT restricts convective oxygen delivery due to contraction-induced increases in IMP.

## Introduction

Physiologically, the critical occluding tension (COT) represents the threshold where contraction induced increases in intramuscular pressure (IMP) exceed intraluminal pressure of the intramuscular vasculature, leading to vascular compression and impairments in muscle perfusion (1–3). The COT has been investigated using isometric contraction models (2, 4, 5) with such muscle actions allowing easier quantification of limb and muscle blood flow (6, 7). Limited research has been conducted examining the COT and its associated physiology in a dynamic resistance exercise model (e.g., contractions involving cyclic concentric and eccentric actions). In a dynamic knee extension resistance exercise model, the COT is estimated to occur between 30-40% 1RM (8) or 40-50% 1RM (9). An oxidative muscle phenotype was positively associated with time to task failure performance at loads ≤30% 1RM, but not loads ≥40% 1RM, supporting the presence of oxygen delivery when exercising below the COT (8). Direct measurement of IMP, limb blood flow, blood pressure, and muscle oxygenation have not been conducted in COT or dynamic resistance exercise models. Simultaneous measures of these variables can provide insight into the physiology of exercise above and below the COT allowing a better understanding of this threshold.

Measurement of IMP requires the placement of an indwelling IMP catheter into the muscle of interest under local anaesthetic (3, 10–12). Given contraction-induced increases in IMP are the physiological basis for the COT, its measurement during exercise is imperative. Microvasculature intraluminal pressure is estimated to be ∼30 mmHg and ∼3-12 mmHg at the arteriolar and venular ends, respectively (13–15). Compartment syndrome research suggests microvascular flow impairments occur when IMP exceeds microvascular intraluminal pressure (16–19). From these studies, it is hypothesized that once contraction-induced increases in IMP exceed venular pressure, perfusion of the muscle will be impaired. Isometric models have found a positive linear relationship exists between relative load and IMP, but individual participant slopes demonstrate substantial variation (3, 10). Therefore, IMP should differ between loads <COT and ≥COT, and specifically, that the IMP occurring at the COT should exceed commonly reported microvascular pressures (13–19).

When exercising to task failure at lower relative loads, IMP increases over time, but at higher relative loads, IMP increases immediately and remains elevated (20). Similar IMP between loading conditions at task failure despite large differences in force requirement (20) suggest motor unit recruitment and firing rate may be important factors explaining IMP. Strong positive relationships between load and normalized electromyography (EMG) amplitude (21, 22), and load and IMP have been previously demonstrated (19, 21), but direct comparisons of normalized EMG amplitude and IMP have yet to be conducted, and such relationships have yet to be quantified in a dynamic resistance exercise model. Muscle action may impact IMP: some studies show lower IMP during maximal eccentric actions of the knee extensors compared to concentric actions (23), but others reported no difference in IMP between maximal concentric, eccentric, and isometric actions of the tibialis anterior and soleus (24). Given dynamic resistance exercise involves cyclical submaximal concentric and eccentric muscle actions, the interaction between IMP and muscle action remains relevant, but equivocal.

Muscle morphological characteristics, as well as placement of the IMP catheter, may also alter IMP when comparing between individuals. Physiological cross-sectional area (PCSA), pennation angle, and/or specific force have been suggested as morphological factors impacting IMP (10) and may also explain between-participant variability in IMP at the same load. The IMP is dependent on catheter depth, with deeper placement eliciting higher pressures (10, 25, 26). Additionally, IMP is also dependent on catheter placement along the muscle length, with proximal placements demonstrating lower pressures compared to distal placements (25, 27). Aiming to place the IMP catheter deep within the central region of the muscle should record the highest pressures compared to superficial and proximal/distal sites. Attempting to match IMP catheter placement sites between participants may be a way to isolate the effects PCSA, pennation angle and specific force have on IMP, but such rigorous placement matching has not been conducted in earlier IMP research (3, 10, 19–21, 23, 27).

Quantification of central and peripheral convective oxygen delivery variables alongside recordings of IMP can help explain the relationship between muscle contraction and perfusion. Dynamic endurance exercise (i.e., alternating cycles of concentric muscle action and relaxation) elicits increases in systolic and diastolic pressures as well as limb specific blood flow up to maximal workloads (28, 29). Early increases in limb blood flow are induced through alterations in the microvascular intraluminal pressure gradient caused by the contraction-relaxation cycle (i.e., the muscle pump) (30), with the relaxation phase following a contraction being associated with large increases in blood flow (31, 32). However, given there are no relaxation periods during dynamic resistance exercise, it is unclear if these findings from endurance exercise apply to resistance exercise models. In an isometric static hold and contract-relax model, loads ≥COT may have impaired microvascular perfusion, as there is minimal leg blood flow occurring during exercise (6). Upon exercise cessation, there may be a larger post-exercise hyperaemic response at loads ≥COT and, thus, the ratio of post-exercise hyperaemic to exercising blood flow would also be increased (33–35). Loads <COT may show elevated exercising blood flow due to limited vascular compression, alongside lower post-exercise hyperaemic responses given the attenuated downstream flow impairment and mechanical compression (33–35).

Near-infrared spectroscopy (NIRS) provides non-invasive insight into the balance between muscle oxygen consumption (mV̇O_2_) and muscle oxygen delivery (mQ̇O_2_), with the difference between these two variables termed muscle oxygenation (SmO_2_). SmO_2_ is shown to be negatively related to exercise intensity (36–38). Previous work found that COT estimates using time to task failure and SmO_2_ were similar, suggesting its ability to reflect exercise states where perfusion is present or absent (8). As femoral artery blood flow does not capture quadriceps perfusion, measuring SmO_2_ in tandem with leg blood flow can provide insight into the relationship between leg blood flow and muscle oxygen status.

This study aimed to examine how mean IMP, leg blood flow, and muscle oxygenation respond to unilateral dynamic resistance knee extension exercise <COT and ≥COT. It was hypothesized that: 1) mean IMP would be positively related to exercise load and normalised EMG, but individual slopes of this relationship would be heterogenous; 2) mean IMP would be highest during concentric compared to eccentric muscle action; 3) exercising leg blood flow and muscle oxygenation would be higher at loads ≤30% 1RM compared to loads ≥40-50% 1RM; 4) post-exercise hyperaemic leg blood flow would be greater at loads ≥40-50% 1RM compared to loads ≤30% 1RM; 5) mean IMP would increase across the lower load task failure bout, but remain elevated across a higher load task failure bout, and no difference in mean IMP would be observed at task failure between bouts.

## Methods

### Ethical Approval

This study was designed and conducted in accordance with the Declaration of Helsinki and approved by the University of British Columbia Research Ethics Board (H25-02203). All participants were informed of the purpose, procedures, and risks of the study prior to providing their written informed consent to participate.

### Participants

Twelve participants (5 female; 7 male) volunteered and consented to participate in the study (Table 1). Participants were between 19 and 40 years of age and able to communicate in English. Participants were ineligible if they had a BMI <18 or >30 kg/m^2^, were a current smoker or user of nicotine devices, had any uncontrolled cardiovascular, metabolic, muscular, joint and/or neurological disorders, had any medical condition affecting their ability to participate in maximal exercise, had been diagnosed or treated for cancer in the past 12 months, or were undergoing any drug therapy that alters skeletal muscle metabolism.

**Table 1.** Participant characteristics.

| **General Information** |  |
|---|---|
| Age (yrs) | 26 (6) |
| Height (m) | 1.73 (0.07) |
| Participants (Female/Male) | 5/12 |
| Body Mass (kg) | 76.9 (13.8) |
| BMI (kg/m<sup>2</sup>) | 25.4 (3.3) |
| **Ultrasound Variables** |  |
| VL MT (cm) | 2.7 (0.4) |
| Thigh ATT (cm) | 0.8 (0.4) |
| VL Sagittal Pennation Angle (°) | 22.1 (3.3) |
| VL Frontal Pennation Angle (°) | 22.8 (2.7) |
| **DXA & Strength** |  |
| Total Body FBFM (kg) | 56.9 (10.2) |
| Thigh FBFM (kg) | 8.3 (1.8) |
| KE 1RM (kg) | 49 (11) |
| Isometric KE MVC (Nm) | 257.1 (62.5) |
| Thigh Specific Force (Nm/kg) | 31.1 (2.3) |
| **Aerobic Performance** |  |
| V̇O<sub>2</sub>peak (ml/kg/min) | 50.9 (8.7) |
| Peak Power Output (W) | 311.8 (59) |
| Relative Peak Power Output (W/kg) | 3.9 (0.6) |

BMI, body mass index; RET, resistance exercise training; VL, vastus lateralis; MT, muscle thickness; ATT, adipose tissue thickness; FBFM, fat-and-bone-free mass; KE, knee extension; 1RM, one-repetition maximum.

### Experimental Design

A within-participant repeated-measures design was used. The study involved three visits: Visits 1 and 2 were the Baseline Testing Visits and Visit 3 was the Intervention Visit. Participants were instructed to consume their usual diet prior to their Intervention Visit, and all participants received the same standardized snack at the same time during their Intervention Visit.

### Baseline Testing Visits Overview

Baseline Testing Visit 1 consisted of basic anthropometry, a full-body dual-energy X-ray absorptiometry (DXA) scan, ultrasound scans of the right vastus lateralis (VL), unilateral knee extension one-repetition maximum (1RM) strength testing, exercise and blood flow occlusion familiarization, and a cycle ergometer V̇O_2_max test. Baseline Testing Visit 2 consisted of a second unilateral knee extension 1RM test to confirm maximal strength, followed by Biodex dynamometry familiarization comprising passive knee extensions and isokinetic and isometric knee extension maximal voluntary contractions (MVCs).

### Anthropometry and Body Composition

Participants wore athletic clothing, removed their shoes, and emptied their pockets before height and body mass were recorded using a stadiometer and scale, respectively. Body composition and fat- and bone-free mass (FBFM) were analyzed using a GE Prodigy iDXA total body scanner (GE Medical Systems, Madison, WI, USA). The DXA scanner was calibrated before each scan using a QA Block Phantom (GE Medical Systems Prodigy, Madison, WI, USA). Custom regions of interest were manually placed to isolate the composition of the right thigh (exercising leg) of each participant as previously described (39, 40).

### Ultrasound Scanning

Ultrasound imaging of the right VL was performed using a GE Logiq i Ultrasound System (GE Medical Systems, Madison, WI, USA), as previously described (40). With participants supine and the leg relaxed, VL length was measured from the anterior superior iliac spine (ASIS) to the superior-lateral corner of the patella. At 50% of VL length, muscle thickness (MT; distance between the superficial and deep aponeuroses), skin thickness, adipose tissue thickness (ATT; distance from the skin to the superficial aspect of the VL), and pennation angle were measured, with values averaged across two scans. Pennation angle was measured in the sagittal plane, with the probe on the anterior aspect of the VL, and in the frontal plane, with the probe on the lateral aspect. Ultrasound variables are presented in Table 1.

### Unilateral Knee Extension Maximal Strength Testing

Unilateral knee extension 1RM was assessed at both Baseline Testing Visits on a plate-loaded machine (C-605 Unilateral Leg Extension, Atlantis Strength, Laval, QC, Canada), and the higher of the two values was used for descriptive purposes and load prescription. The 1RM was defined as the heaviest load lifted through the full range of motion (ROM), from ∼90° of knee flexion to ∼0-5° of knee flexion, independent of contraction velocity. Seat depth was adjusted to align the knee with the loading arm’s axis of rotation, and the shin pad ROM adjustment was set to its deepest setting for all participants. Following a warm-up of 5-10 submaximal repetitions and 3-6 repetitions at a slightly heavier load, participants performed four to six progressively heavier single repetitions until their 1RM was reached. Load was increased by 0.6-4.5 kg after each successful attempt or reduced by 0.6-1.1 kg if needed to confirm the 1RM, with 2 min of rest between attempts.

### Exercise Familiarization

Following 1RM testing, participants were familiarized with the Intervention Visit exercise by completing ten repetitions at 30% 1RM and three repetitions at 60% 1RM. Tempo (1.5 s concentric, 1.5 s eccentric) was paced using a custom auditory and visual metronome, and ROM matched that used during 1RM testing.

### Cycle Ergometry Maximal Oxygen Uptake (V̇O_2_peak)

Whole-body peak oxygen consumption (V̇O_2_peak) and peak power output were determined using a ramp incremental test to volitional failure on an electronically controlled cycle ergometer (Monark LC6, Monark Sports and Medical, Vansbro, Sweden). Following 2-3 min of resting baseline data collection and a 3-5 min warm-up at 20-100 W (selected based on participant body size and training history), power output increased by 20 W/min until volitional failure or until cadence could not be maintained above 60 rpm. Gas exchange data were collected using a metabolic cart (Quark RMR, COSMED, Rome, Italy) with OMNIA software (COSMED). V̇O_2_peak was taken as the highest 30 s average of V̇O_2_, and peak power output as the highest power output achieved.

### Dynamometry Familiarization and Testing

Dynamometry familiarization and testing was performed on a Biodex dynamometer (Biodex Medical Systems, Inc., Shirley, NY, USA; software version 5.3.06). The lateral epicondyle of the right femur was aligned with the dynamometer’s axis of rotation, and the shin pad was positioned ∼2.5 cm proximal to the lateral malleolus. ROM was set from each participant’s maximal achievable knee flexion (∼110°) to ∼5-10° of knee flexion. For both the familiarization and testing participants first completed two sets of 15 passive knee extensions while remaining fully relaxed. They then performed three sets of isokinetic knee extensions at 45°/s, with each repetition comprising a concentric contraction immediately followed by an eccentric contraction. Following the isokinetic efforts participants performed three sets of 5 s isometric contractions at 60° of knee flexion. For both contraction types, participants were instructed to “kick into the pad as hard and as fast as possible” on a “go” signal, with 2 min of rest between sets. During the familiarization session the three sets per test were performed at ∼75%, ∼85%, and ∼95-100% effort, and during testing each set was performed with maximal effort.

### Intervention Visit Overview

The Intervention Visit consisted of 13 unilateral knee extension bouts (Table 2), with Biodex dynamometer bouts performed first, followed by dynamic resistance exercise bouts. Bouts 1-3 (dynamometry) followed the familiarization procedures described above. Bouts 4-11 involved dynamic isotonic knee extension resistance exercise against varying relative and absolute loads performed for a submaximal number of repetitions. Bouts 12 and 13 were performed to volitional failure, defined as the participant electing to stop, or two consecutive repetitions falling outside the prescribed ROM bounds or tempo (1.5 s concentric, 1.5 s eccentric). Before, during, and after each dynamic bout (e.g., bouts 4-13), VL IMP, femoral artery blood flow, blood pressure, VL muscle oxygenation, VM EMG, and kicking arm ROM were recorded continuously (Figure 1). IMP, ROM, and blood pressure were recorded in LabChart (Supplementary Figure 1), whereas muscle oxygenation, EMG, and blood flow were recorded on independent systems and time-synchronized post-collection, where required. Participants consumed a standardized snack (one bottle of juice and one granola bar) during the 20 min rest period between Bouts 11 and 12.

**Figure 1.**
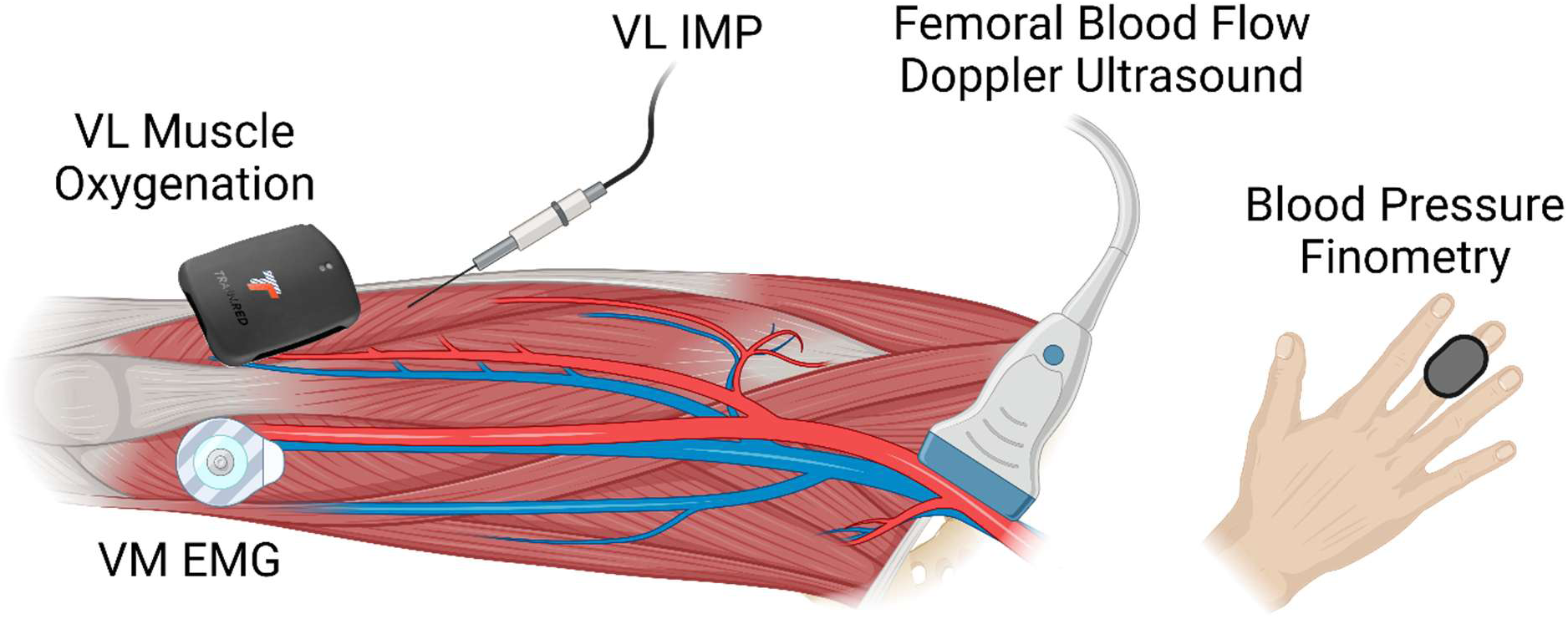
Visual of study measures setup. Vastus lateralis (VL) intramuscular pressure (IMP) and muscle oxygenation were measured alongside vastus medialis (VM) electromyography (EMG), common femoral artery blood flow using Doppler ultrasound, and reconstructed blood pressure using finger finometry. Created in https://BioRender.com

**Table 2.** Intervention visit exercise bout details.

| Order | Exercise Bout | Equipment | Sets | Repetitions | Rest (min) |
|---|---|---|---|---|---|
| 1 | Passive knee extension | Biodex dynamometer | 2 | 15 | 2 |
| 2 | Maximal isokinetic knee extension (concentric and eccentric) |  | 3 | 1 | 2 |
| 3 | Maximal isometric knee extension |  | 3 | 1 | 2 |
| 4 | 20% 1RM | Plate-loaded knee extension | 1 | 10 | 5 |
| 5 | 30% 1RM |  | 1 | 8 | 5 |
| 6 | 40% 1RM |  | 1 | 6 | 5 |
| 7 | 50% 1RM |  | 1 | 6 | 5 |
| 8 | 60% 1RM |  | 1 | 6 | 5 |
| 9 | 10 kg |  | 1 | 10 | 5 |
| 10 | 15 kg |  | 1 | 8 | 5 |
| 11 | 20 kg |  | 1 | 6 | 5 |
| Standardized snack |  |  |  |  |  |
| 12 | 60% 1RM | Plate-loaded knee extension | 1 | Maximum | 20 |
| 13 | 20% 1RM |  | 1 | Maximum | - |

1RM, one-repetition maximum; kg; kilograms

### IMP Catheter Pre-Placement Calculations

IMP catheter placement is depicted in Figure 2. The Catheter Insertion Point equation standardized the insertion site at 65% of the distance from the ASIS to the superior-lateral corner of the patella, so that the catheter tip would lie in the middle portion of the muscle belly, and the Length of Catheter Inserted equation standardized tip depth at 60% of VL MT. Inputs for these equations were VL length, MT, ATT, and sagittal plane pennation angle.

**Figure 2.**
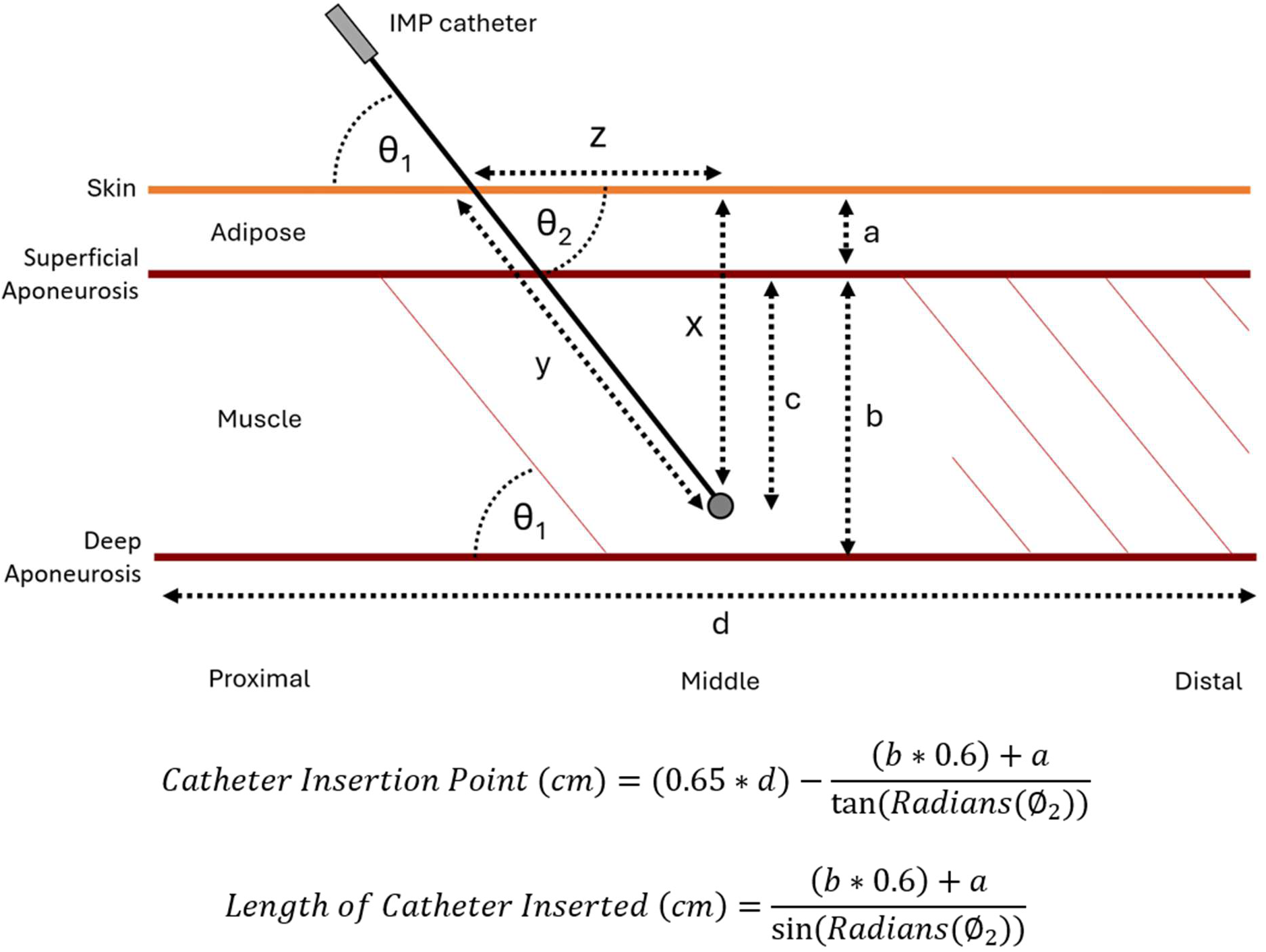
Insertion variables and equations. Ultrasound derived variables include: *θ* as the muscle pennation/insertion angle (°), *x* as the distance from the skin to the catheter tip (cm), *y* as the length of the catheter inserted into the muscle (cm), *z* as the distance from the insertion point in the skin to the catheter tip (cm), *a* as the adipose/skin thickness (cm), and *b* as the muscle thickness (cm), *d* as the length of the muscle (cm). The IMP catheter tip depth *c* is selected by the research team as a desired percentage of muscle thickness.

### IMP Catheter Placement Procedure

Before each visit, participant-specific Catheter Insertion Point and Length of Catheter Inserted equations were completed using ultrasound data from Baseline Testing Visit 1. Participants lay supine with their leg relaxed, and the Catheter Insertion Point was marked on the VL. Following local anaesthesia of the skin and fascia (2% Xylocaine with epinephrine; 1:100,000) and ultrasound-guided infiltration of deeper tissues along the planned insertion path, the site was prepared with Betadine Solution (10% povidone-iodine). Under sterile conditions and ultrasound guidance, a physician inserted an introducer needle with a peel-away Teflon cannula (ON-Q T-Peel Introducer Needle, Avanos Medical Inc., Alpharetta, GA, USA; Supplementary Figure 2) at the Catheter Insertion Point, matching the sagittal and frontal plane pennation angles as closely as possible so that the needle ran parallel to the muscle fibers. The needle was advanced until its tip reached ∼60% of MT and was then removed, leaving the cannula in place. A sterile IMP catheter (Mikro-Cath Pressure Catheter, Millar Instruments, Houston, TX, USA), pre-marked so that its tip would sit level with the cannula tip, was inserted through the cannula to this mark. The cannula was then withdrawn while the catheter was held in place, and torn away along its perforations. The catheter was secured to the thigh with porous medical tape beneath a sterile dressing (Figure 3, Panel A), and its tip was located by ultrasonography to quantify catheter depth within the muscle, using the same imaging procedures as Baseline Testing Visit 1 (Figure 3, Panel B).

**Figure 3.**
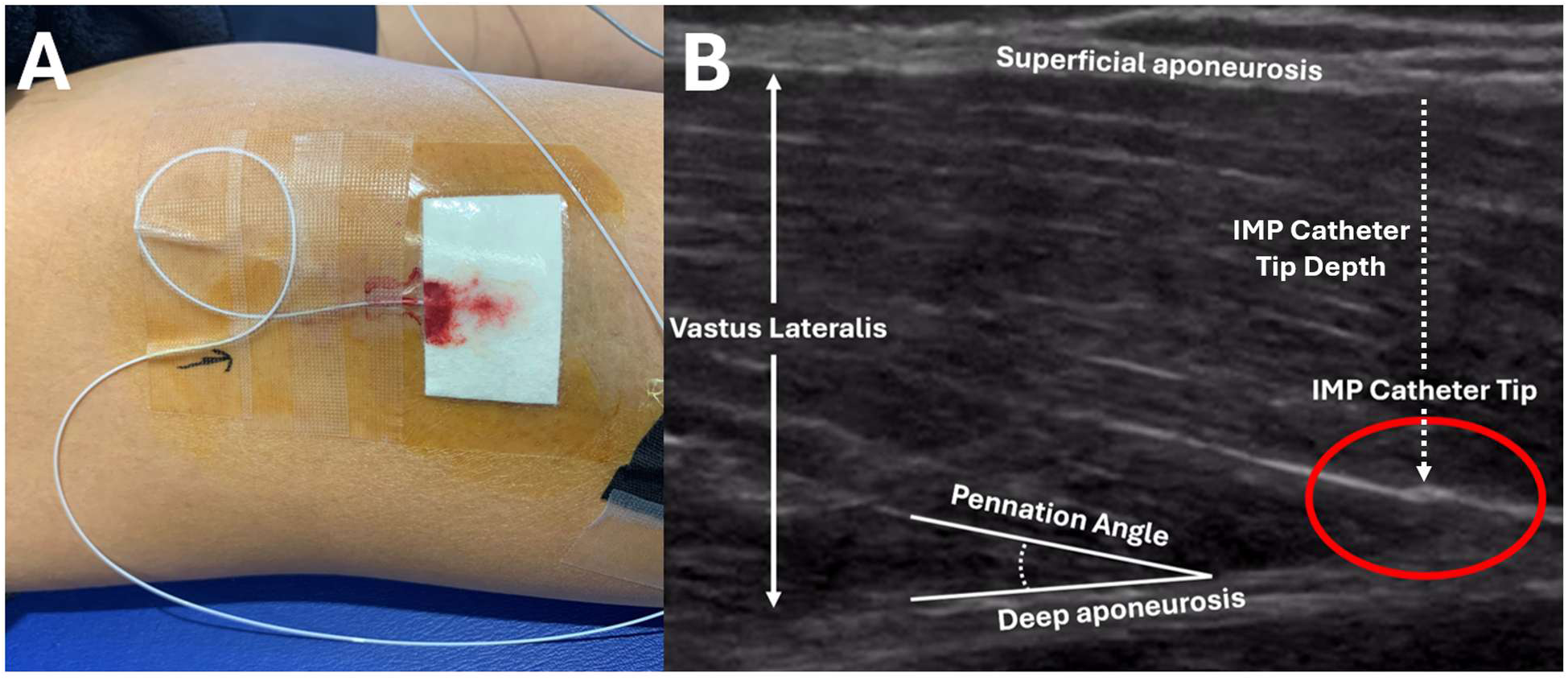
Sample image of IMP catheter placement. Panel A depicts the placed IMP catheter secured via Tegaderm pad and porous medical tape; a loop in the catheter wire was used to reduce catheter motion once placed. Panel B shows an ultrasound image of the IMP catheter tip within the VL; the IMP catheter wire can be seen to be nearly parallel with the pennation angle.

### IMP Catheter Data Recording

The IMP catheter was connected in series to an interface cable (PEC-10D Cable, Millar Instruments, Houston, TX, USA), a pressure control unit (PCU-2000, Millar Instruments, Houston, TX, USA), and a PowerLab system (PowerLab; ADInstruments, Colorado Springs, CO, USA) for LabChart integration. IMP data were recorded at 1000 Hz and exported at 10 Hz for subsequent analysis. Catheters were calibrated following removal while remaining connected to the PowerLab system. Calibration procedures are provided as supplementary materials (Supplementary Figure 3).

### Data Analysis Overview

For all relative- and absolute-load non-failure bouts, the first six repetitions were used for analysis of all measures. For the relative load failure bouts, each bout was split into tertiles based on repetitions completed, and only the repetitions in the first and third tertiles were used for IMP, blood pressure, and EMG analyses. Additionally, only the first and last three repetitions of each relative non-failure bout were used for leg blood flow analysis.

### IMP Data Analysis

IMP data were analysed in RStudio (41) after smoothing with a fourth-order low-pass Butterworth filter (1 Hz cut-off). Resting IMP was calculated as the mean IMP over 30 s while participants were seated on the dynamometer or knee extension machine. For dynamometry bouts, maximum IMP was determined for each passive knee extension and averaged across both passive bouts. Isokinetic repetitions (4 s) were divided by time into 2 s concentric and eccentric phases. For isokinetic and isometric MVCs, rolling 0.5 s averages of torque and IMP were calculated, and the highest 0.5 s torque average was taken as peak torque, along with the corresponding 0.5 s IMP average. For knee extension machine bouts, individual repetitions were identified from joint ROM data, and repetitions not reaching the required ROM were excluded. Concentric and eccentric phases were defined by ascending and descending joint angles, respectively, and each was divided at the midpoint of the repetition’s ROM into bottom and top portions, yielding four phases: bottom concentric, top concentric, top eccentric, and bottom eccentric (Supplementary Figure 4). For non-failure bouts, mean IMP was calculated across the entire bout, and maximum, minimum, and phase-specific mean IMP were calculated per repetition and averaged across the bout. For task failure bouts, mean IMP and per-repetition maximum and minimum IMP were averaged within each tertile.

### Leg Blood Flow

Common femoral artery diameter and blood velocity were measured by a trained sonographer (KC) using a 10 MHz probe and a GE Logiq i Ultrasound System (GE Medical Systems, Madison, WI, USA) for 30 s before, throughout, and for 30 s after each bout. Arterial diameter and anterograde, retrograde, and net (anterograde minus retrograde) velocities were analysed using Cardiovascular Suite 4 (Quipu, Pisa, Italy), and blood flow was calculated as cross-sectional area multiplied by velocity and by 60 to express flow per minute, then averaged into 1 s bins. Resting and post-exercise hyperaemic flows were taken as the mean of the 20 s immediately before and after each bout, respectively. For non-failure relative-load, and absolute-load bouts, exercising flow was taken as the mean across the first six repetitions. For task failure bouts, exercising flow was calculated in two ways: as the mean of the first and last three repetitions, and as the mean within the first (T1) and third (T3) tertiles after dividing each bout into tertiles based on repetitions completed. Exercising and post-exercise hyperaemic flows were expressed relative to each bout’s pre-bout resting flow.

### Blood Pressure

Beat-by-beat blood pressure was measured at the left hand using height-corrected finger photoplethysmography (Finometer, Finapres Medical Systems BV, Arnhem, The Netherlands) and calibrated against manual blood pressure measurements taken from the right arm at heart level every five bouts. Pressure traces were analysed using a custom R script to obtain mean systolic and diastolic pressure and cardiac cycle count. For non-failure relative-load, and absolute-load bouts, these variables were averaged across the first six repetitions. For task failure bouts, they were calculated across the first and last three repetitions and within T1 and T3, with cardiac cycles expressed per repetition for the tertile analysis.

### Muscle Oxygenation

Muscle oxygenation of the VL was recorded continuously using a commercially available NIRS device (Train.Red FYRE 2.0, Train.Red, Elst, The Netherlands). Data were processed in a custom application (42), where they were resampled at 10 Hz, smoothed with a fourth-order low-pass Butterworth filter (0.2 Hz cut-off), and normalised to a 0-100 scale using the minimum and maximum values in each participant’s data file. Individual bouts were then manually identified and extracted for analysis using a custom R script. Within each bout, the 3 s preceding the lowest SmO_2_ value were averaged to yield a 3 s minimum SmO_2_ value, which was used for statistical analysis.

### EMG

Surface EMG of the vastus medialis (VM) was recorded using a portable Biometrics EMG system (DataLog MWX8 with SX230-1000 sensors; Biometrics Ltd., Newport, UK). Electrodes were aligned along the centre of the VM muscle belly ∼4-5 cm above the superior border of the patella, with a ground strap (R506) placed at the lateral malleolus of the non-exercising leg, and signals were zeroed with the participant at rest. EMG was sampled at 1000 Hz with an internal 10-450 Hz band-pass filter and analysed in MATLAB using a custom script (GY), with no additional filtering. Repetitions were identified using a moving standard deviation window (200 samples), and peak amplitude was extracted from a 500-sample moving RMS envelope for each repetition. Peak RMS values were averaged within each isometric MVC, non-failure relative-load, and absolute-load, and within tertiles for task failure bouts, then normalised to the mean peak RMS achieved during the MVC trials.

### Joint Range of Motion

Joint ROM was measured using a custom inertial measurement unit (IMU; developed by AC) interfaced with PowerLab and converted to degrees in LabChart using a two-point calibration. Visual bounds on the ROM channel provided real-time feedback to participants and researchers to ensure that full ROM was achieved, and ROM data were synchronized with IMP to enable position-specific IMP analyses.

### Work and Work Rate

Work was calculated from IMU-derived ROM data using a custom R script. Repetitions were identified by peak and valley detection of joint ROM, and concentric work for each repetition was calculated from the participant-specific load and the fixed moment arm of the knee extension machine (0.35 m), then summed. As the number of repetitions differed between relative and absolute loads, total work was calculated over the first six repetitions of non-failure relative, and absolute load bouts, and across repetitions within T1 and T3 of task failure bouts. Work rate was calculated as work completed divided by bout duration.

### Statistical Analysis

Data are presented as mean (SD) unless otherwise stated (43). Analyses were performed in SPSS (v29.0.2.0; IBM Corp., Armonk, NY, USA) with significance set at α≤0.05. All comparisons used linear mixed-effects models with participant as a random effect, and extreme outliers identified in SPSS were removed.

Fixed effects were muscle action (concentric, eccentric, isometric) for dynamometry-derived IMP and torque; relative load (20%, 30%, 40%, 50%, and 60% 1RM) or absolute load (10, 15, and 20 kg) for non-failure IMP, leg blood flow, change in femoral artery diameter, blood pressure, SmO_2_, and EMG. Within-repetition IMP for non-failure bouts was analysed with contraction phase (bottom concentric, top concentric, top eccentric, bottom eccentric) and relative load as fixed effects, and in a reduced model with load grouped relative to the COT (20-30% vs. 40-60% 1RM). Task failure IMP, leg blood flow, blood pressure, SmO_2_, and EMG were analysed with load (20% vs. 60% 1RM) and tertile (T1 vs. T3) as fixed effects.

Repeated-measures correlations coefficients (44, 45) were calculated between relative load and normalised EMG, between mean IMP and relative or absolute load, and between mean IMP and normalised EMG. Individual-participant linear regressions were also fitted for the mean IMP relationships, and model fits are reported. Pearson correlations were calculated between minimum SmO_2_ and 1RM strength at each relative and absolute load; between mean IMP and 1RM strength, thigh FBFM, specific force, pennation angle, and absolute and relative catheter tip depth; and between mean IMP at 20 kg and 1RM strength, thigh FBFM, and specific force.

## Results

### IMP Catheter Placement

There was no effect of depth type for relative IMP catheter depth when comparing between calculated and actual depths (Calculated = 60 (0.0) %MT; Actual = 57.6 (11.0) %MT; p=0.47). However, there was an effect of depth type for absolute IMP catheter depth with the actual absolute depth being shallower than the calculated depth (Calculated = 1.7 (0.2) cm; Actual = 1.3 (0.39) cm; p<0.01). Together these results indicate that the correct relative depth was achieved during IMP catheter placement, but differences between ultrasound image plane and site may explain the absolute catheter depth differences.

### Biodex Dynamometry

Passive repetition IMP data are descriptively presented in Figure 4 alongside the resting IMP values and mean IMP achieved during each relative load. Peak torque was highest during the eccentric action, compared with both the concentric and isometric maximal actions (p<0.05), and peak torque during the isometric action was higher than during the concentric action (p<0.05). Comparison of IMP at peak torque showed moderate-to-large effects between muscle actions, but significance was not reached (p>0.05). Similarly, the main effect of muscle action was not significant for IMP per unit torque (p>0.05), but effect size estimates suggest the isometric action produced higher IMP per unit torque. Data are presented in Table 3.

**Figure 4.**
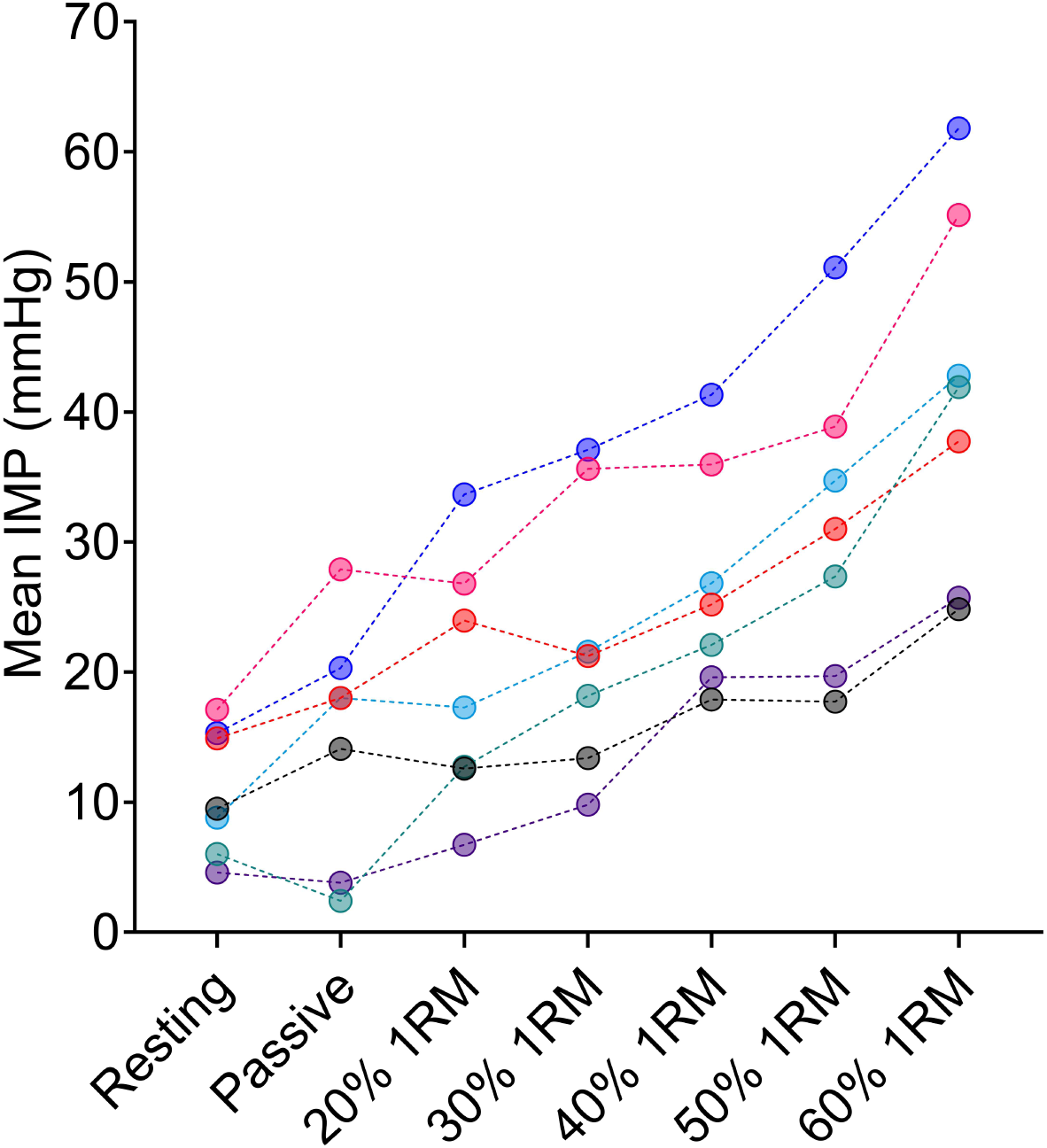
Individual participant descriptive IMP data. Descriptive plot of each participants resting IMP, maximum IMP recorded during dynamometry passive knee extension repetitions, and mean IMP across each relative load bout.

**Table 3.** Dynamometry torque and IMP values.

|  | **Maximal Concentric** | **Maximal Eccentric** | **Maximal Isometric** | **p** | **Con vs Ecc Cohen’s d** | **Con vs Iso Cohen’s d** | **Ecc vs Iso Cohen’s d** |
|---|---|---|---|---|---|---|---|
| Peak Torque (Nm) | 225.3 (57.7) | 309.1 (92.2)<sup>*</sup> | 257.1 (62.5)<sup>*</sup> | <0.01 | - | - | - |
| IMP at Peak Torque (mmHg) | 41.8 (23.0) | 64.4 (39.8) | 91.1 (64.7) | 0.09 | d=0.7 | d=1.0 | d=0.5 |
| IMP per unit Torque (mmHg/Nm) | 0.17 (0.11) | 0.18 (0.10) | 0.31 (0.21) | 0.06 | d=0.0 | d=0.9 | d=0.8 |

Con, concentric; Ecc, eccentric; Iso, isometric; IMP, intramuscular pressure. <sup>\*</sup>Significantly different from Con

### Knee Extension

#### Work, Work Rate, and Repetitions Completed

##### Non-Failure Relative Loads

A significant main effect of relative load was found for work completed across the first six repetitions (p<0.01), and work rate (p<0.01). All loads showed significantly different work across the first six repetitions (p<0.05) except for 30% 1RM and 40% 1RM, which were similar (p=0.57) (20% 1RM = 0.56 (0.12) kJ; 30% 1RM = 0.69 (0.18) kJ; 40% 1RM = 0.70 (0.17) kJ; 50% 1RM = 0.86 (0.20) kJ; 60% 1RM = 1.1 (0.23) kJ). Work rate was significantly different between all tested loads (p<0.01). Work rate data are presented in Figure 5, Panel A.

**Figure 5.**
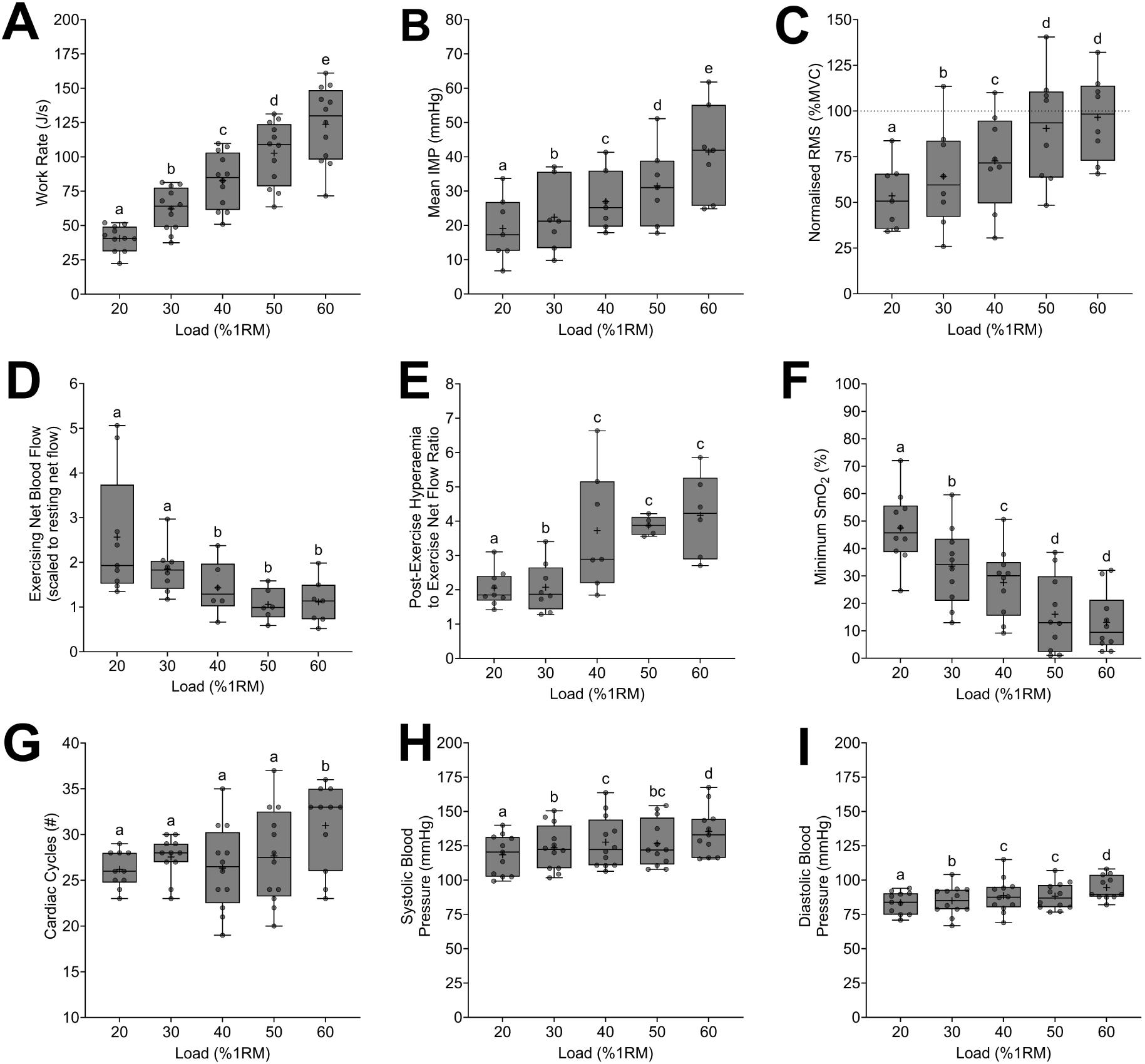
Non-failure relative load data. Between load comparisons for work rate (A), mean IMP (B), EMG (C), exercising net blood flow (D), post-exercise hyperamia:exercise net blood flow ratio (E), minimum SmO_2_ (F), cardiac cycle count (G), mean systolic blood pressure (H), and mean diastolic blood pressure (I) are presented. Data are presented to show the median (horizontal line), interquartile range (boxes), minimum and maximum (bars), and mean (+). Different letters are significantly different from one another (p<0.05).

##### Task Failure Relative Loads

There was a significant main effect of task failure relative load on work (p<0.01) and work rate (p<0.01). More work was performed in the 20% 1RM condition than in the 60% 1RM condition (20% 1RM = 4.6 (1.4) kJ; 60% 1RM = 1.9 (0.7) kJ; p<0.01). Work rate was higher in the 60% 1RM condition than the 20% 1RM condition (20% 1RM = 40.8 (8.0) J/s; 60% 1RM = 131.2 (50.8) J/s; p<0.01). Data are presented in Figure 6, Panel A. Participants completed 80 (36) repetitions in the 20% 1RM task failure condition, and 10 (2) repetitions in the 60% 1RM task failure condition. The total time spent exercising was 4.0 (1.8) min, and 0.5 (0.1) min for the 20% and 60% 1RM task failure conditions, respectively.

**Figure 6.**
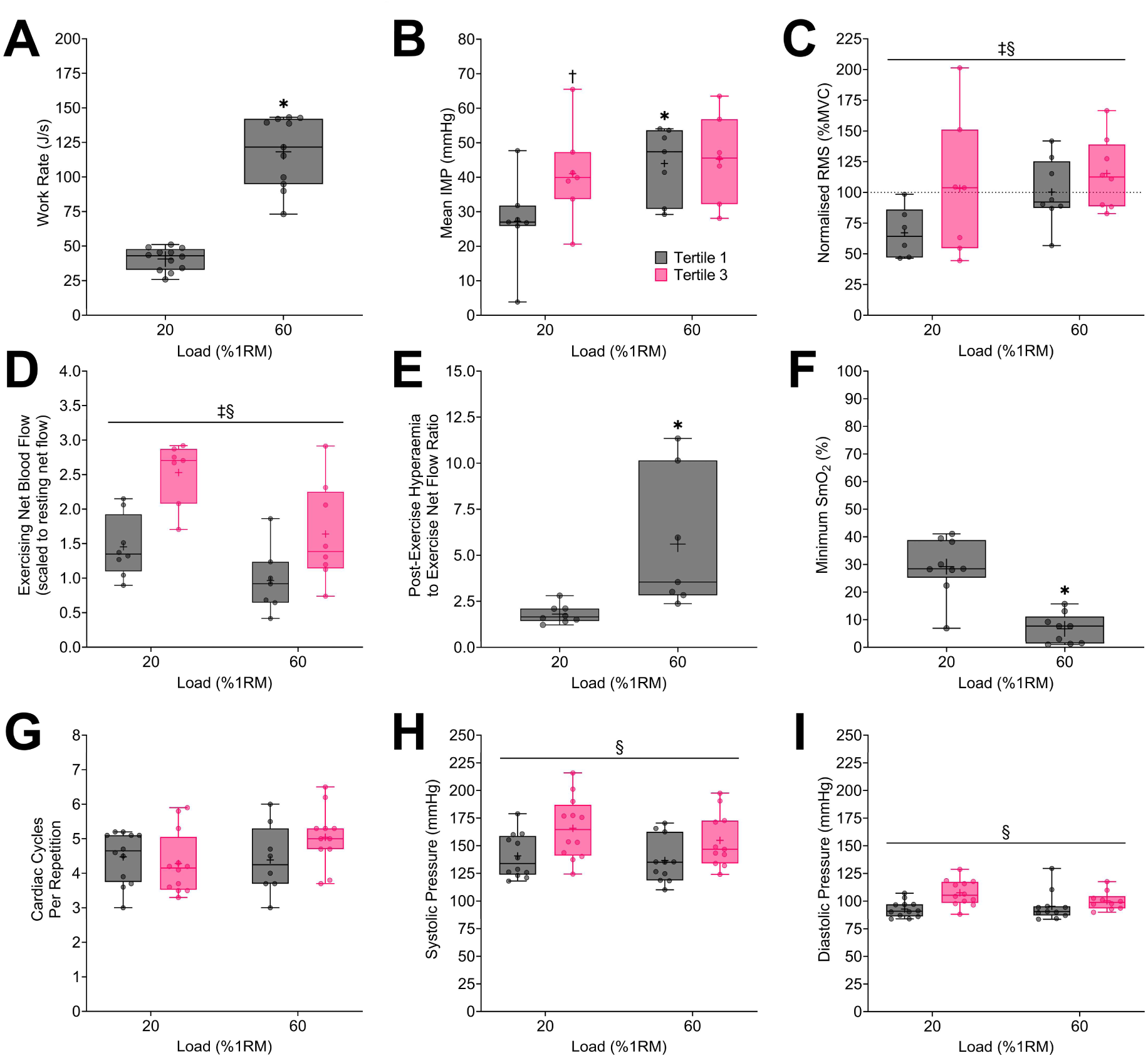
Task failure relative load data. Between load comparisons for work rate (A), mean IMP (B), EMG (C), exercising net blood flow (D), post-exercise hyperamia:exercise net blood flow ratio (E), minimum SmO_2_ (F), cardiac cycle count (G), mean systolic blood pressure (H), and mean diastolic blood pressure (I) are presented. Data are presented to show the median (horizontal line), interquartile range (boxes), minimum and maximum (bars), and mean (+). Different letters are significantly different from each other (p<0.05). * Significant difference between relative loads within a tertile (p<0.05). † Significant difference between tertiles within a relative load (p<0.05). ‡ Significant main effect of relative load (p<0.05). § Significant main effect of tertile (p<0.05).

#### Intramuscular Pressure

Out of 12 participants, seven presented with usable IMP data (Supplementary Figure 5, Panel A), and five presented with unusable IMP data. Four participants did not display a physiologically representative change in IMP during exercise which we defined as no response in the IMP pressure values during contraction that otherwise should have induced a change (Supplementary Figure 5, Panel B). One participant had an inverted IMP signal (Supplementary Figure 5, Panel C). These five participants did not have their IMP data included in analysis.

##### Non-Failure Relative Loads

A significant main effect of relative load was found for maximum IMP (p<0.01), mean IMP (p<0.01), and minimum IMP (p=0.03). Maximum IMP was different between all loads (p<0.05) except for 20% and 30% 1RM did not differ from each other (p>0.05) (20% 1RM = 34.4 (25.1) mmHg; 30% 1RM = 39.8 (19.1) mmHg; 40% 1RM = 50.2 (16.2) mmHg; 50% 1RM = 55.8 (19.2) mmHg; 60% 1RM = 71.3 (20.9) mmHg). Mean IMP was different between all loads (p<0.05). Minimum IMP was higher at 60% 1RM than at all other loads (p<0.05), with all other loads not differing from one another (p>0.05) (20% 1RM = 10.6 (7.0) mmHg; 30% 1RM = 11.6 (9.4) mmHg; 40% 1RM = 11.9 (7.9) mmHg; 50% 1RM = 15.5 (11.9) mmHg; 60% 1RM = 22.5 (12.5) mmHg). Mean IMP data are presented in Figure 5, Panel B.

##### Task Failure Relative Loads

A significant interaction was found for relative load × tertile for mean IMP (p=0.02). The 60% 1RM condition showed a higher mean IMP than the 20% 1RM condition at T1 (p<0.01), but there was no difference between loads at T3 (p<0.01). Mean IMP significantly increased from T1 to T3 in the 20% 1RM condition (p<0.01) but did not differ between tertiles for the 60% 1RM condition (p=0.71). A significant main effect of task failure relative load was found for maximum IMP (p=0.02). Collapsed across tertiles, maximum IMP was greater at 60% 1RM than at 20% 1RM (20% 1RM = 51.2 (30.5) mmHg; 60% 1RM = 70.2 (27.4) mmHg; p=0.02). No significant main effects or interactions were found for task failure minimum IMP. Mean IMP data are presented in Figure 7, Panel B.

##### Contraction Phase

Significant main effects of contraction phase (p<0.01) and load (p<0.01) were found, but the interaction was non-significant (p=0.20). Collapsed across load, each contraction phase was significantly different from all other contraction phases (all p<0.05). Data are presented in Figure 8, Panel A.

**Figure 8.**
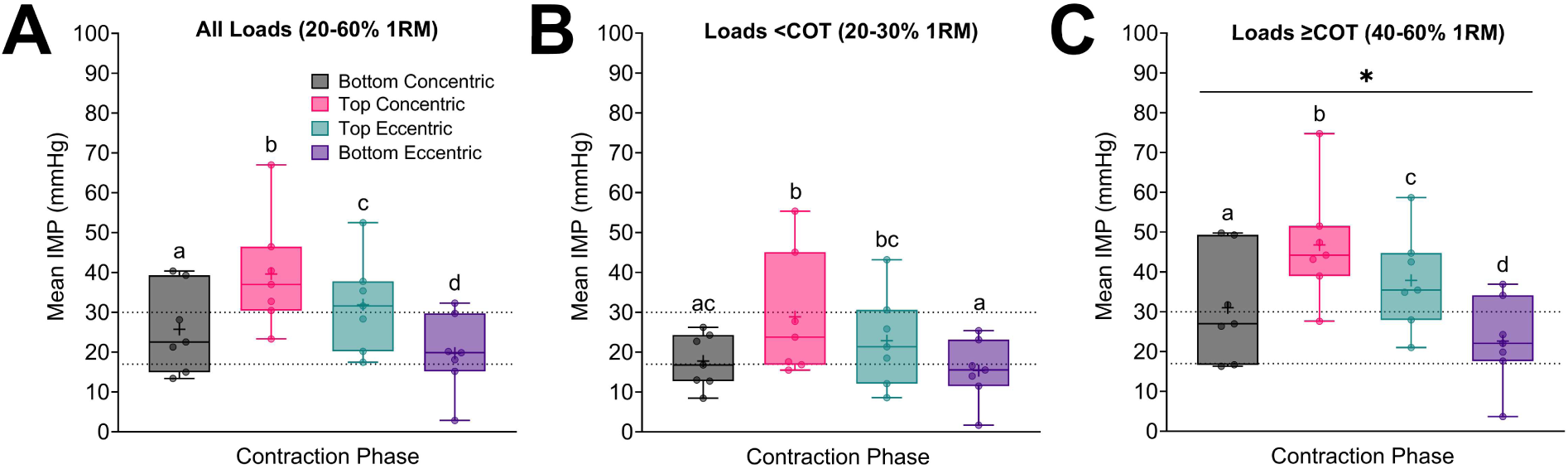
Mean IMP across the knee extension contraction phases. Panel A represents the mean IMP per repetition phase collapsed across all loads. Panel B reports the mean IMP across the repetition phases for only loads <COT. Panel B shows the mean IMP across the repetition ROM for only loads ≥COT. Data are presented to show the median (horizontal line), interquartile range (boxes), minimum and maximum (bars), and mean (+). Horizontal dashed lines represent estimation of microvascular compression (17 mmHg) and microvascular occlusion (30 mmHg). Different letters are significantly different from one another (p<0.05). * Significant difference between loads <COT and ≥COT for each repetition phase.

From the reduced model, a significant interaction of load × contraction phase was found (p=0.02). Within each contraction phase, mean IMP was higher in the ≥COT load condition than in the <COT load condition (Figure 8, Panels B and C). In the <COT load condition, bottom concentric was similar to bottom eccentric and top eccentric, top concentric was similar to top eccentric, and bottom eccentric was only similar to bottom concentric (Figure 8, Panel B). All contraction phases were different from each other within the ≥COT load condition (all p<0.05) (Figure 8, Panel C).

##### Relationship between Load and Mean IMP

A significant repeated-measures correlation coefficient of (r_rm_=0.92; 95% CI [0.86, 0.95]; p<0.01) was found for the relationship between relative load and mean IMP, and between absolute load and mean IMP (r_rm_=0.90; 95% CI [0.83, 0.94]; p<0.01). Data are presented in Figure 9.

**Figure 9.**
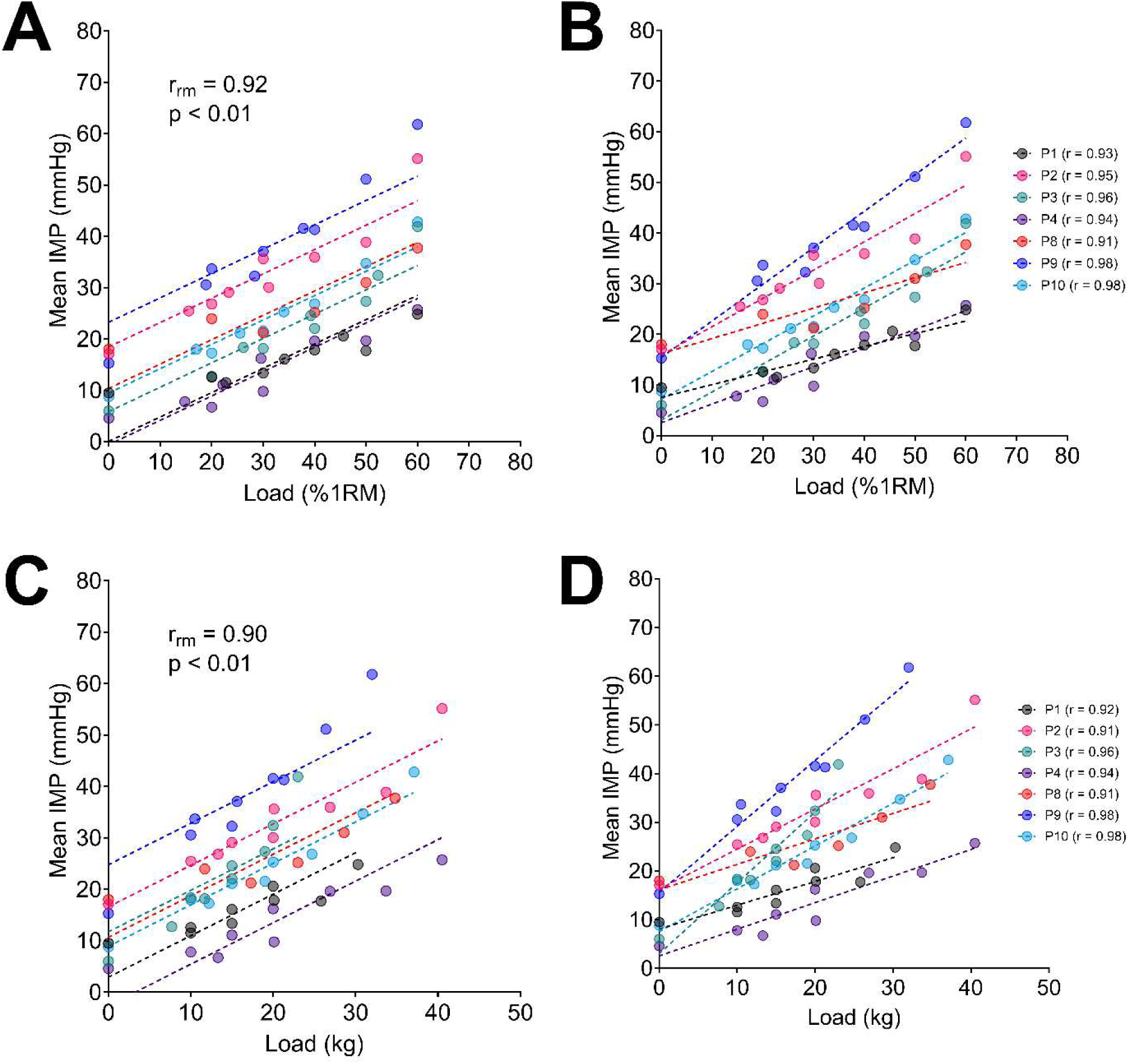
Relationship between mean IMP and load. The left panels show repeated measures correlation coefficient analysis between mean IMP and relative (A) and absolute (C) loading. The right panels show individual participant slopes for the relationship between mean IMP and relative (B) and absolute (D) loading.

##### Relationship between mean IMP and descriptive variables

No significant relationships were found between mean IMP (collapsed across relative loads) and knee extensor 1RM strength (r=0.02; p=0.76), thigh FBFM (r=-0.03; p=0.94), knee extensor specific force (r=0.39; p=0.39), pennation angle (r=0.74; p=0.06), absolute catheter tip depth (r=0.29; p=0.53), or relative catheter tip depth (r=-0.04; p=0.93). Data are presented in Figure 10. Further, no significant relationships were found between mean IMP at 20 kg and 1RM strength (r=-0.31; p=0.55), thigh FBFM (r=-0.49; p=0.32), or specific force (r=0.43; p=0.47). Data are presented in Figure 11.

**Figure 10.**
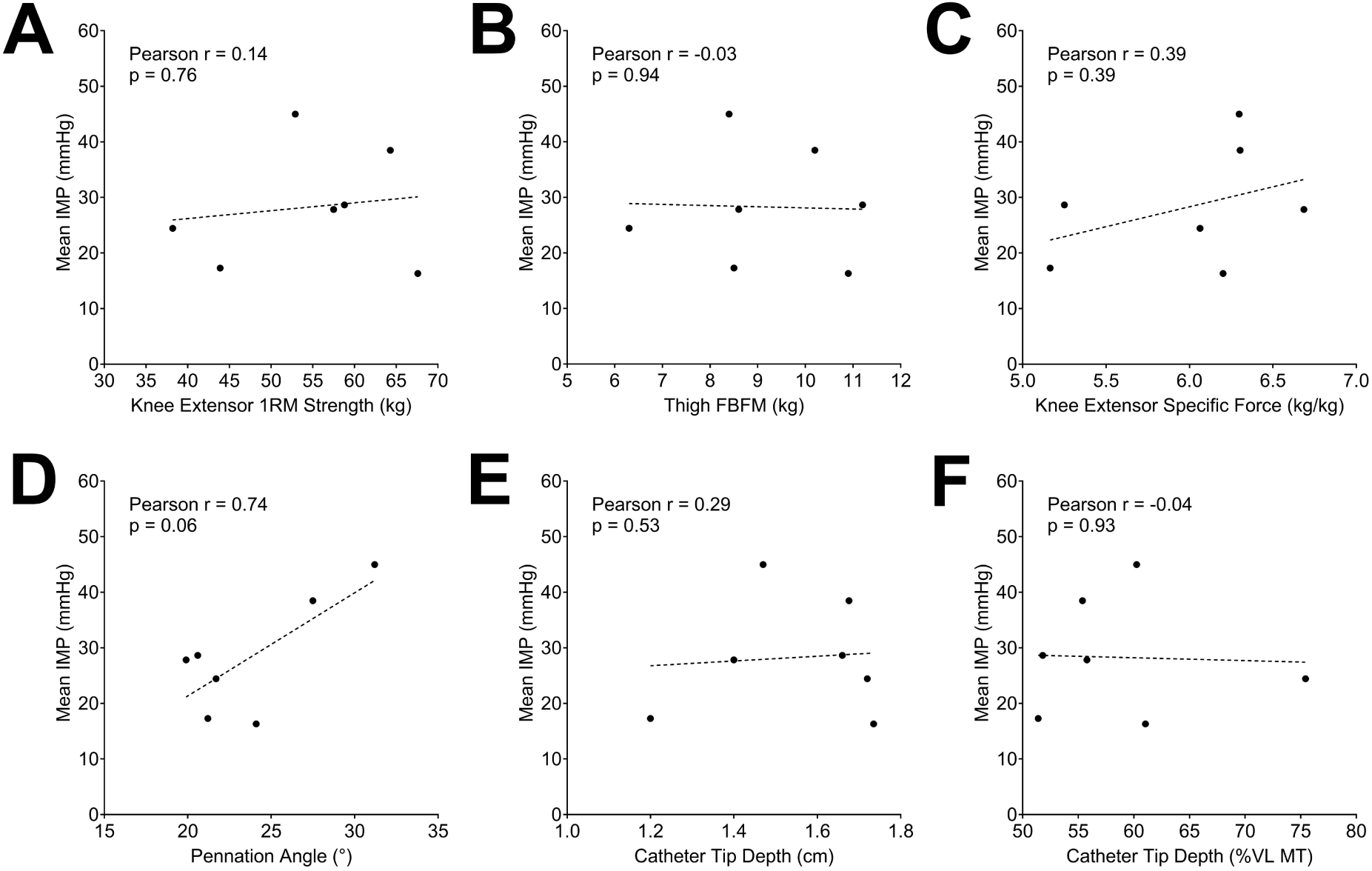
Correlation between relative load mean IMP and descriptive variables. Mean IMP collapsed across relative load (20-60% 1RM) is plotted against knee extensor 1RM strength (A), thigh fat-and-bone-free mass (FBFM) (B), knee extensor specific force (C), pennation angle (D), absolute catheter tip depth (E), and relative catheter tip depth (F). Pearson correlation coefficients are provided for each graph alongside p-values.

**Figure 11.**
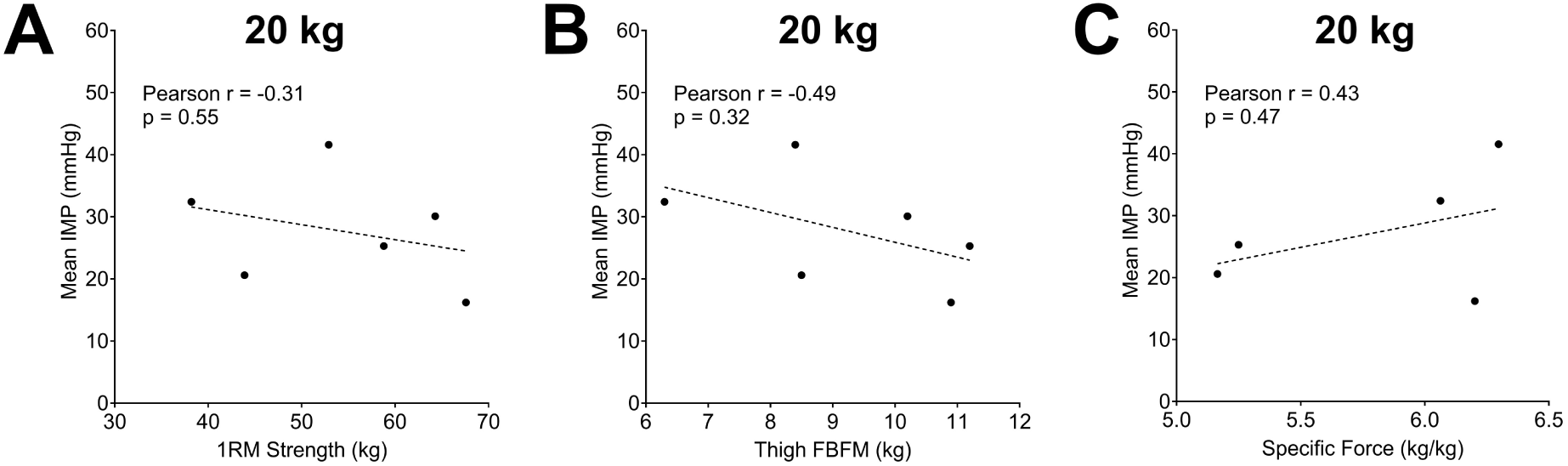
Correlation between 20 kg mean IMP and muscle strength, size, and specific force. Mean IMP across the 20 kg bout is plotted against knee extensor maximal strength (A), and thigh FBFM (B). Pearson correlation coefficients are provided for each graph alongside p-values.

#### EMG

##### Non-Failure Relative Loads

A significant main effect of relative load was observed for normalised RMS (p<0.01). Normalised RMS at 50% and 60% 1RM did not differ (p=0.28), but all other relative load comparisons were significantly different (p<0.05). Data are presented in Figure 5, Panel C.

##### Failure Relative Loads

Significant main effects of relative load (p<0.01) and tertile (p<0.01) were found, but the interaction was non-significant (p=0.19). Collapsed across tertile, normalised RMS was greater at 60% 1RM than at 20% 1RM. Collapsed across relative load, normalised RMS was higher at T3 than at T1. Data are presented in Figure 7, Panel C.

##### Relationship between mean IMP and normalised EMG

Significant repeated-measures correlation coefficients were found for the relationship between relative load and normalised RMS (r_rm_=0.90; 95% CI [0.81, 0.95]; p<0.01), pooled mean IMP from all relative-load and absolute-load bouts and normalised EMG (r_rm_=0.78; 95% CI [0.62, 0.89]; p<0.01), and mean IMP and normalised RMS using pooled tertile data from the 20% 1RM and 60% 1RM task failure bouts (r_rm_=0.82; 95% CI [0.54, 0.93]; p<0.01). Data are presented in Figure 12.

**Figure 12.**
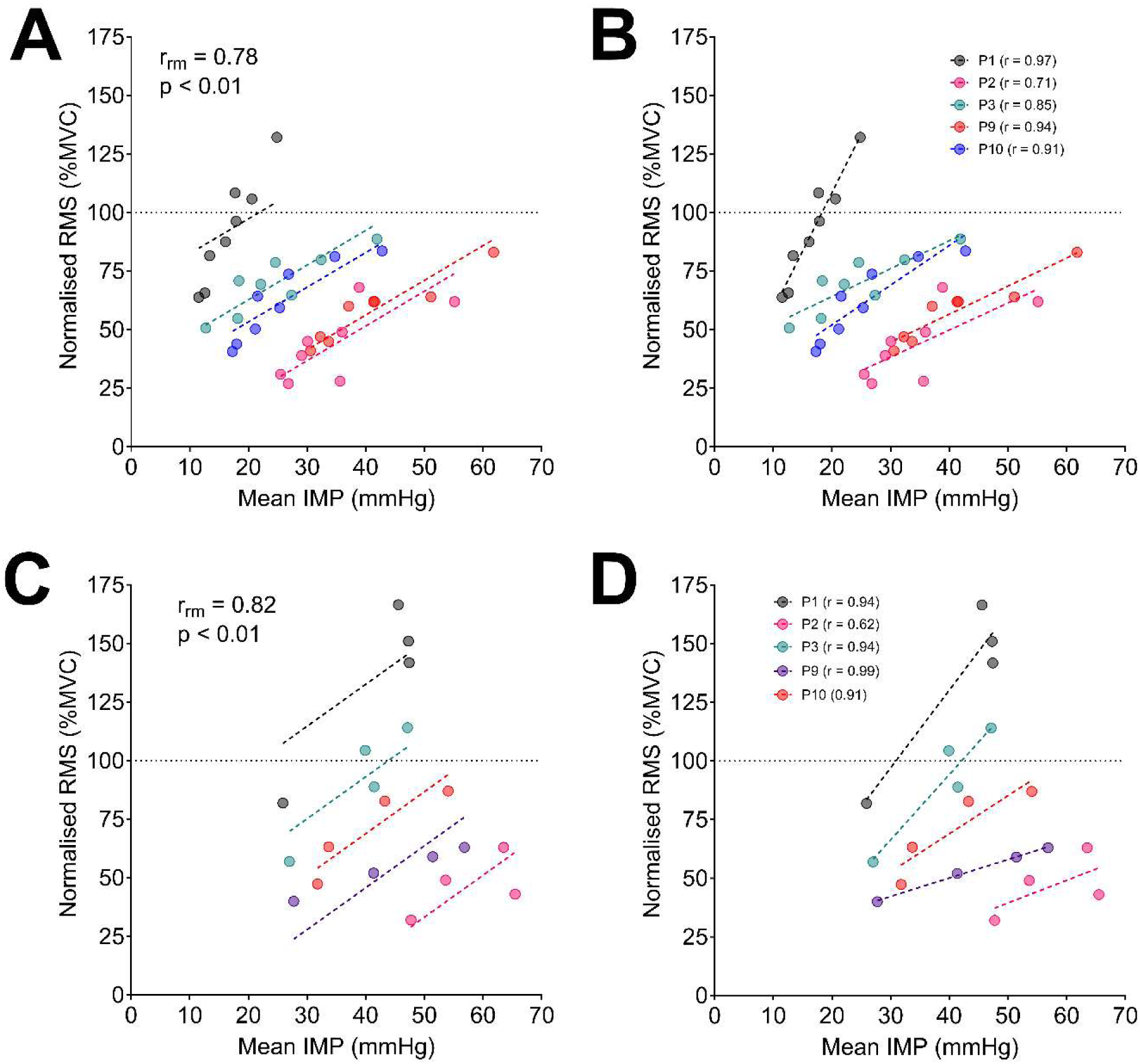
Relationship between mean IMP and normalised EMG. The left panels show repeated measures correlations for pooled relative and absolute non-failure loads (A) and pooled tertile data from the 20% 1RM and 60% 1RM task failure bouts (C). The right panels show individual participant slopes and their corresponding R^2^ values for pooled relative and absolute non-failure loads (B), and pooled tertile data from the 20% 1RM and 60% 1RM task failure bouts (D).

#### Leg Blood Flow

Out of 12 participants, 2 participants did not have femoral blood flow measured due to scheduling limitations.

##### Non-Failure Relative Loads

No effect of relative load was found for the absolute change in femoral artery diameter from pre-exercise to during exercise (20% 1RM = 0.02 (0.09) mm; 30% 1RM = -0.01 (0.06) mm; 40% 1RM = -0.05 (0.11) mm; 50% 1RM = 0.00 (0.03) mm; 60% 1RM = -0.04 (0.14) mm; p=0.52).

No effect of relative load was found for resting net blood flow (20% 1RM = 0.40 (0.26) L/min; 30% 1RM = 0.32 (0.15) L/min; 40% 1RM = 0.35 (0.08) L/min; 50% 1RM = 038 (0.17) L/min; 60% 1RM = 0.42 (0.18) L/min; p>0.05).

A significant main effect of relative load was found for net exercising leg blood flow scaled to rest (p<0.01). There was no difference between 20% 1RM and 30% 1RM (p>0.05), nor were 40% 1RM, 50% 1RM and 60% 1RM different from one another (p>0.05). However, both 20% 1RM and 30% 1RM were significantly different from to 40% 1RM, 50% 1RM, and 60% 1RM (p<0.05). Data are presented in Figure 5, Panel D.

No effect of relative load was found for post-exercise hyperaemic net blood flow scaled to rest (20% 1RM = 6.2 (2.8) times rest; 30% 1RM = 5.5 (2.0) times rest; 40% 1RM = 5.5 (1.8) times rest; 50% 1RM = 4.7 (1.7) times rest; 60% 1RM = 4.3 (1.2) times rest; p=0.37).

A significant main effect of relative load was found for the post-exercise hyperaemic:exercise net blood flow ratio (p<0.01). There was no difference between 20% 1RM and 30% 1RM (p=0.78), nor were 40% 1RM, 50% 1RM and 60% 1RM different from one another (p>0.05). However, both 20% 1RM and 30% 1RM were significantly different compared from 40% 1RM, 50% 1RM, and 60% 1RM (p<0.05). Data are presented in Figure 5, Panel E.

##### Failure Relative Loads

Examination of absolute time-matched comparisons (first versus last three repetitions) showed significant main effects of relative load (p<0.01) and time (p<0.01), but there was no interaction effect (p=0.27). Collapsed across time, exercising net blood flow scaled to rest was greater in the 20% 1RM condition than in the 60% 1RM condition (p<0.01). Collapsed across relative loads, exercising net blood flow scaled to rest was greater across the last three repetitions than across the first three repetitions (p<0.01). Data are presented in Figure 7, Panel D.

Examination of relative time-matched comparisons (T1 versus T3) showed significant main effects of relative load (p<0.01) and tertile (p=0.01). Collapsed across relative load, exercising net blood flow scaled to rest was at T3 than at T1 (T1 = 1.7 (1.0) times rest; T3 = 2.2 (1.0) times rest; p<0.01). Collapsed across tertile, exercising net blood flow scaled to rest was higher in the 20% 1RM condition than in the 60% 1RM condition (20% 1RM = 2.6 (0.8) times rest; 60% 1RM = 1.3 (0.7) times rest; p=0.01).

#### Blood Pressure

##### Non-Failure Relative Loads

A significant main effect of relative load was found for cardiac cycle count (p<0.01), systolic blood pressure (p<0.01), and diastolic blood pressure (p<0.01). Cardiac cycle count was significantly greater at 60% 1RM than at all other loads (p<0.05), and all other loads did not differ from each other (p>0.05). Systolic blood pressure was significantly lower at 20% 1RM than at all other loads (p<0.01) and significantly higher at 60% 1RM than at all other loads (p<0.01). Systolic blood pressure at 30% 1RM was different from all other loads (p<0.05) except 50% 1RM (p=0.17), and at 40% 1RM it was different from all other loads (p<0.05) except 50% 1RM (p=0.61). Systolic blood pressure at 50% 1RM was higher than at 20% 1RM (p<0.01), no different from 30% 1RM and 40% 1RM (p>0.05), and lower than at 60% 1RM (p<0.01). Diastolic blood pressure was similar between 40% 1RM and 50% 1RM (p=0.77), but all other relative load comparisons were significantly different (p<0.05). Data are presented in Figure 5, Panels Panel G, H, and I.

##### Failure Relative Loads

Examination of absolute time-matched comparisons (first versus last three repetitions) showed a significant main effect of time for cardiac cycle count (p<0.01) and systolic blood pressure (p<0.01), while a significant relative load × time interaction for diastolic blood pressure was found (p=0.02). Collapsed across relative loads, cardiac cycle count, and systolic blood pressure increased from the first to the last three repetitions (p<0.01). Diastolic blood pressure was greater at 60% 1RM than at 20% 1RM across the first three repetitions (p<0.01), increased in both relative loads between the first and last three repetitions (p<0.01), and was no different between relative loads across the last three repetitions (p=0.34). Data are presented in Figure 7, Panels G, H, and I.

Examination of relative time-matched comparisons (T1 versus T3) showed a significant main effect of tertile for both systolic and diastolic blood pressure (p<0.01), but no significant main effects or interactions were present for cardiac cycles per repetition (p>0.05). Collapsed across relative loads, systolic blood pressure was at T3 than at T1 (T1 = 139.7 (21.4) mmHg; T3 = 161.1 (28.0) mmHg; p<0.01), as was diastolic blood pressure (T1 = 94.9 (11.1) mmHg; T3 = 107.1 (15.1) mmHg; p<0.01), but cardiac cycles per repetition were similar (T1 = 4.1 (1.2) cycles/repetition; T3 = 4.6 (0.9) cycles/repetition; p>0.05).

#### Muscle Oxygenation

##### Non-Failure Relative Loads

A significant main effect of relative load was found for minimum 3-second SmO_2_ (p<0.01). All loads were significantly different from one another (p<0.01), except for 50% 1RM and 60% 1RM which did not differ (p=0.16). Data are presented in Figure 5, Panel F.

##### Failure Relative Loads

A significant main effect of relative load was found for minimum 3-second SmO_2_ (p<0.01). Minimum 3-second SmO_2_ was higher in the 20% 1RM task failure condition than in the 60% 1RM task failure condition (p<0.01). Data are presented in Figure 7, Panel F.

##### Relationship between SmO_2_ and 1RM Strength

No correlation between minimum 3-second SmO_2_ and 1RM strength was observed at 20% 1RM (r=0.00; p=0.94), or 30% 1RM (r=0.24; p=0.50). Weak non-significant correlations were seen at 40% 1RM (r=0.33; p=0.36), 50% 1RM (r=0.50; p=0.14), and 60% 1RM (r=0.58; p=0.08). A moderate significant correlation was observed at 10 kg (r=080; p<0.01), with large significant correlations observed at 15 kg (r= 0.97; p<0.01), and 20 kg (r=0.94; p<0.01). Data are presented in Figure 13.

**Figure 13.**
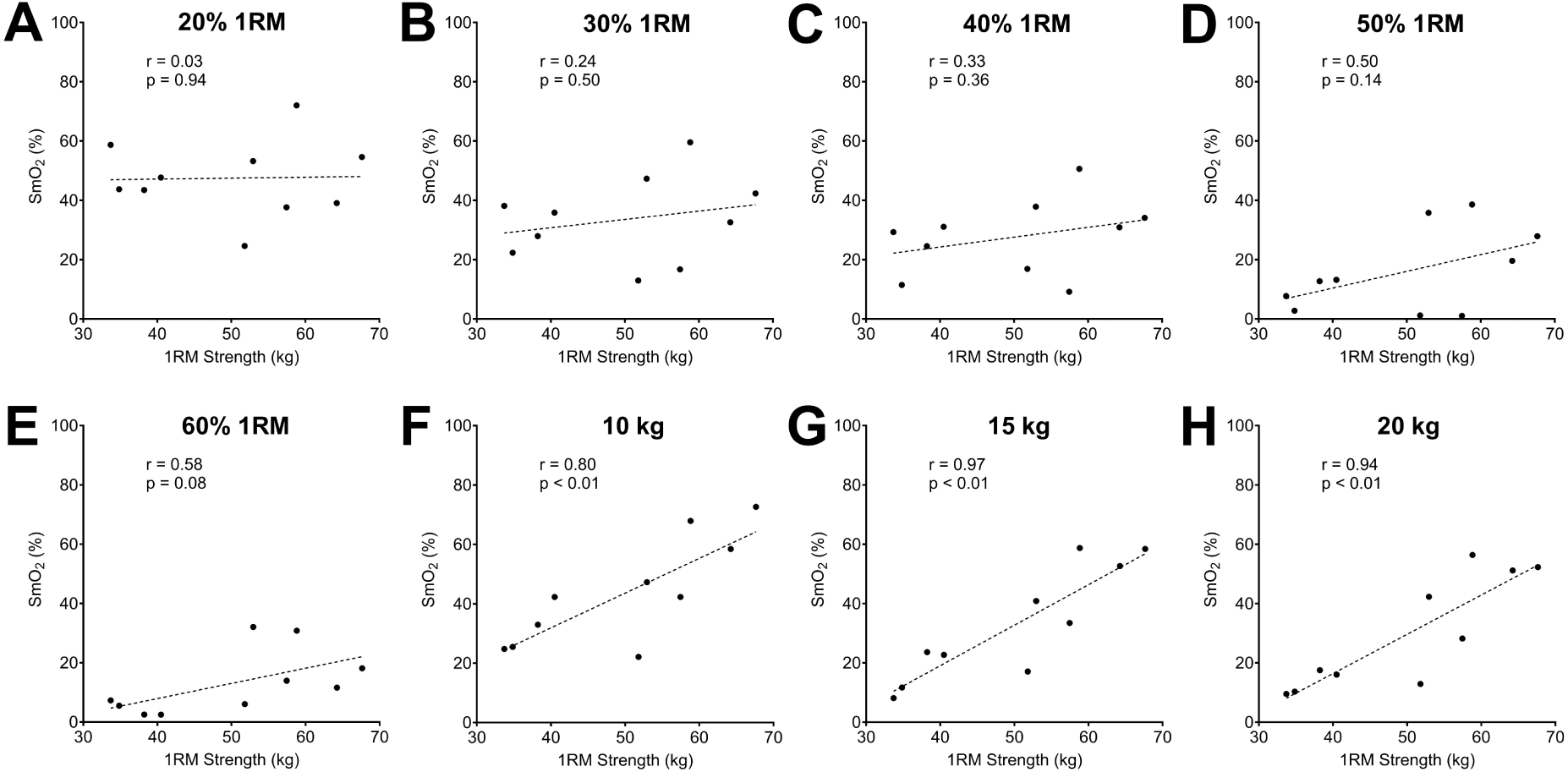
Relationship between minimum 3-second SmO_2_ and 1RM strength. Panels A-E present the SmO_2_ and 1RM strength relationship for relative loads, and Panels F-H present the within absolute load SmO_2_ and 1RM strength relationship.

## Discussion

This study used integrative physiological measures to investigate how dynamic resistance exercise load influences convective oxygen delivery. First, the present findings align with previous work from our group identifying the dynamic unilateral knee extension COT to be between 30 and 40% 1RM (8). Exercising blood flow and muscle oxygenation were reduced at loads ≥40% 1RM compared to loads ≤30% 1RM despite blood pressure increasing with load. Further, the site of this convective oxygen transport limitation was likely downstream of the femoral artery, at the level of the intramuscular microvasculature. Net exercising blood flow did not reduce during exercise at any load nor did it reach zero suggestive of occlusion, but contraction-induced mean IMP increased with load and likely exceeded microvascular intraluminal pressure at loads ≥40% 1RM. Second, when exercising to volitional failure, mean IMP increased over time when working against a lower load but increased immediately and stayed elevated over time when exercising with higher loads. Further, there was no difference in mean IMP at task failure between the lower and higher loads. This result suggests that, although IMP is positively related to relative load, the number of active muscle fibers and their firing rate may be the principal underlying mechanism governing contraction-induced increases in IMP, given similar IMP values at task failure despite different force requirements.

Earlier work using a dynamic knee extension resistance exercise model suggests the lower bound of the COT phase transition occurs at loads >30% 1RM and the upper bound occurs at ∼40-50% 1RM (8, 9, 46). These loads align with the current study’s results, as exercising net blood flow was substantially lower at loads ≥40% 1RM than at loads ≤30%1RM. Our blood flow findings are also consistent with work showing reduced leg blood flow during the contraction phase of isometric knee extension exercise using a contract-relax model when loads were ≥30% MVC (6). Given that contraction may have had little effect on the common femoral artery itself, the reduced exercising blood flow at loads ≥40% 1RM likely stems from downstream microvascular impairment. During exercise, blood flow is greatest during muscle relaxation (32) and initially increases via the muscle pump effect (i.e., contraction compressing vessels and relaxation opening vessels, which increases flow), and mechanically triggered vasodilation (30, 31). With no true relaxation period present during dynamic resistance exercise the microvasculature may remain occluded across the full ROM. Mean IMP was higher at each contraction phase for loads ≥COT (i.e., 40-60% 1RM) than for loads <COT (i.e., 20-30% 1RM). Not only were mean IMP values lower at loads <COT, but the bottom portions of the concentric and eccentric phases may also represent periods in which perfusion can occur due to very low contraction-induced IMP. Future work combining similar IMP measures on a plate-loaded resistance exercise machine with better time-synchronized limb blood flow and muscle oxygenation measures could test this hypothesis.

Post-exercise hyperaemia was similar across relative loads despite differences in work completed and work rate. Post-exercise hyperaemia would be expected to be positively related to contraction load, work completed, or work rate, which does not align with our data (30, 47, 48). The lower loads (20-30% 1RM) in our study demonstrated post-exercise hyperaemia comparable to that of the higher loads (40-60% 1RM) despite lower contraction intensities, lower total work completed, and a lower work rate across the bouts. However, bout duration was longer for the lower loads by ∼6-9 s due to differences in repetitions completed which may increase the oxygen debt of the exercising muscle, thereby increasing post-exercise hyperaemia. At loads ≥40% 1RM, the post-exercise hyperaemia is likely caused by greater occlusion of the microvasculature and subsequent higher mechanical and metabolic stimuli for vasodilation upon removal of the mechanical compression (49–51). This mechanical compression would prevent increases in exercising flow, as these vessels remain collapsed, but would instead result in a large post-exercise hyperaemic response once contraction ceases. The above mechanisms, while rooted in previous work (30, 47, 52, 53), remain speculative, given that the exercise bouts were not matched for duration and thus the effect of bout duration cannot be separated from the overall findings.

The ratio of post-exercise hyperaemia to exercising blood flow can provide insight into the effect of contraction intensity on microvascular flow as blood flow is tightly matched to metabolic demand (48, 54, 55). The ratio of post-exercise hyperaemia to exercising blood flow showed a breakpoint between 30% and 40% 1RM, which aligns with the previously established COT for this specific exercise model (8). Post-exercise hyperaemic flow was greater than exercising flow across all loads but was ∼2-fold higher at 20-30% 1RM, compared with ∼4-fold higher from 40-60% 1RM. This ratio can be thought of as an index of the magnitude of contraction-induced flow impairment, analogous to damming a river. At 20-30% 1RM, blood flow is impaired by contraction but is still maintained, with demand for blood increasing as contractions continue. Exercising flow to the leg increases, and, once contraction ceases, flow increases further in the post-exercise state, but the magnitude of this change relative to exercise is modest. In contrast, as exercising blood flow barely changed from rest at 40-60% 1RM, it is proposed that the downstream vasculature was nearly completely blocked. Once contraction ceases, a large percentage of microvasculature undergoes vasodilation, elicited from mechanical and metabolic stimuli, alongside an increase in the microvascular pressure gradient leading to increases in post-exercise hyperaemic flow. Given that post-exercise hyperaemic flow was similar across all loads, whereas 20-30% 1RM produced increases in exercising flow and 40-60% 1RM produced minimal increases, the ratio of post-exercise hyperaemic flow to exercising flow is inherently larger for the 40-60% 1RM loads than for the 20-30% 1RM loads.

A single quadriceps muscle group makes up ∼11.5% of total lower-body muscle volume (56), and, as such, the cardiovascular demand during unilateral knee extension exercise is lower than during cycling or running. Despite this, systolic and diastolic blood pressure increased with relative load, indicating a response from the cardiovascular system. Starting at the lower bound of the COT (i.e., 30% 1RM), blood pressure began to increase, which may be a sympathetically mediated response to overcome the mechanical compression occurring at the muscle, consistent with previous knee extensor findings (57). This interpretation is consistent with the exercising blood flow results: minimal flow increases at loads ≥COT (i.e., ≥40% 1RM) due to occlusion (mechanical), combined with increased oxygen demand (metabolic), would ultimately lead to an increased pressor response. Further, at these higher loads, the widespread local vasodilation paired with the increased pressor response would support a large increase in post-exercise hyperaemic flow. Additionally, exercising blood flow was lower at 40-60% 1RM than at 20-30% 1RM despite the positive association between blood pressure and relative load. Together, these findings suggest that the occlusive nature of higher-load contractions leads to a central increase in pressure to perfuse the working muscle. Despite these increased blood pressures, limb blood flow was unable to increase suggesting that peripheral occlusion may override central blood pressure increases. f

Mean IMP was positively related to relative load at the group level, similar to previous reports using isometric contraction models (10). Novel to this study were the identification of a similar relationship during dynamic contractions and the individual-level linear correlations, which demonstrated between-participant differences in resting IMP (y-intercept) and the IMP response to load (slope) (Figure 9). Participants with similar slopes did not necessarily have similar resting IMP values, and vice versa, demonstrating the independence of these two factors when comparing load-IMP relationships between participants. For descriptive purposes, resting IMP, peak IMP from passive knee extension repetitions, and mean IMP across each relative load bout were plotted (Figure 4). This provides further nuance to the IMP response to contraction as passive knee extensions, which alter muscle shape in the absence of force generation, and produced a general increase in IMP above rest in most participants. Overall, some component of the contraction-induced increase in IMP must arise from muscle shape change, although the force generation component of IMP appears to be the largest contributor to its change.

Another novel aspect of this study was the measurement of IMP across different portions of the ROM, capturing the effect of muscle action and torque on IMP. Collapsed across loads, the highest IMP was observed at the top of the concentric phase, and the lowest at the bottom of the eccentric phase. This aligns with the torque demands of the plate-loaded knee extension machine used in this study (Supplementary Figure 6), as the top of the ROM is where torque is highest, and the bottom is where it is lowest. Further, concentric actions are associated with greater radial expansion of the muscle belly (58) a require a greater number of active fibers at a fixed load than eccentric actions. Mean IMP was also found to be positively related to normalised EMG at the group and individual levels. Previous work has shown a similar relationship (3, 21) and has therefore related contraction-induced IMP generation to the total number and firing rate of motor units. This theory aligns with earlier IMP work identifying factors relevant to contraction-induced IMP generation, in which specific force was listed as a key variable. It should be noted that EMG was recorded from the VM, while IMP was measured within the VL. This discrepancy in muscles measured was due to limited space on the VL for all devices (e.g., IMP, NIRS, EMG) to be administered simultaneously, and so VM EMG was chosen, as the VM produces an EMG pattern similar to that of the VL during knee extension exercise (59).

The effect that exercising to task failure at higher and lower loads has on IMP has been examined previously in an isometric model (20) and was the basis for our inclusion of such bouts in this study. Our experiment built upon this work by using a dynamic resistance exercise model while measuring IMP alongside cardiovascular variables and muscle electrical activity to provide an integrative assessment of LME performance. During the first 33% of the task failure bouts, IMP was higher in the heavier-load condition, but during the last 33%, there was no difference in IMP between loads. IMP appeared to increase immediately to its maximum in the heavier-load condition but to increase over time in the lower-load condition. This pairing of results may relate to the motor unit recruitment patterns at each load. Normalised EMG amplitude data showed that quadriceps muscle electrical activity was higher in the heavier-load than in the lower-load condition, and in the last third than in the first third of exercise, but did not follow the same pattern as mean IMP. However, the relationship between mean IMP and normalised EMG amplitude at the group level was strong (r_rm_=0.82) and was maintained at the individual level (r=0.71-0.99). Motor unit recruitment and firing rate would plausibly be related to IMP, given the strong associations between IMP and load, and between EMG and load found in this study and in previous work (3, 10, 21, 60).

Greater muscle electrical activity is associated with increased motor unit recruitment and firing rate (61), both of which allow force to be increased (62) or maintained (63). This relationship likely explains why IMP is so tightly related to both relative load and normalised EMG amplitude in the present study. During a sustained contraction, initially recruited muscle fibers fatigue, and more fibers must be recruited to maintain force; this is consistent with the EMG findings from the present study’s task failure bouts. However, mean IMP was immediately elevated and maintained across the entire higher-load task failure bout but started lower and increased over time in the lower-load task failure bout, with both reaching similar mean IMP values at the end of exercise. This similarity in end-of-bout mean IMP between loads appears to contradict the established relationship between IMP and relative load, but it may underscore another aspect of the IMP-contraction relationship. At the end of a task failure bout, it is assumed that near complete recruitment of motor-units will have occurred regardless of the load or exercise duration (64). IMP may then be more closely related to the total number of motor units recruited and their activation rate, with the maximum achievable IMP occurring at the point where an individual is maximally recruiting and activating all of their available muscle fibers. This theory is consistent with the non-failure findings, given that muscle fiber recruitment during a submaximal task is strongly related to the required force output, but it only partially supports the task failure results, as EMG amplitude was larger overall in the 60% 1RM than in the 20% 1RM condition, and larger at T3 than at T1. Previous work has shown lower EMG values at task failure in lower-compared to higher-load conditions, but muscle fibre glycogen depletion is similar between conditions suggesting similar total fibre use across the duration (64). There is a lack of research examining EMG and IMP simultaneously, so some speculation is required regarding how the present findings fit into contraction physiology. When exercising to task failure using lower loads, individual fibres will fatigue leading to less force generated per fibre but a time-related increase in total fibres producing force. In contrast, higher-load task failure will immediately recruit larger motor units at a higher firing rate, and due to the shorter contraction duration, potentially fewer fibres will undergo fatigue leading to many fibres being recruited and producing higher forces per fibre. As IMP was shown to reach similar values at task failure between lower- and higher-load bouts in the present study, there is some discrepancy between the EMG and IMP findings. A simple proposition would be that IMP is directly related to the number of fibres recruited and their individual force outputs, but at task failure the force per fibre is likely lower at the end of lower-load bouts. It could also be assumed that a 1RM test would produce the highest achievable IMP due to maximal motor unit recruitment being present in the absence of individual fibre fatigue; in the present study, IMP was not recorded during a 1RM attempt, so this proposition remains speculative. To summarize, this study provides novel insight into the relationship between exercise load, EMG, and IMP showing strong relationships between these variables. However, precise examination of the total number of fibres recruited and the force per fibre across loads and task failure conditions would provide much needed context for the proposed theories stemming from our findings.

### Limitations

Several limitations of the present study warrant discussion. First, the IMP traces were not physiologically responsive in 5 of 12 participants. In one participant, the signal appeared inverted, as confirmed by joint ROM data, and in four participants, the IMP response to contraction was far below that expected at the prescribed loads (Supplementary Figure 5, Panels B and C). The cause of these errors is unclear. In our study, catheters were not connected to PowerLab until insertion was complete; connecting them beforehand (while keeping the inserted portion sterile) would allow faulty catheters to be identified and placement to be adjusted if recorded pressures appeared physiologically implausible. Second, IMP was measured in a single region of the VL. Catheter depth and longitudinal position clearly affect recorded IMP within the same muscle contracting at the same intensity (10, 25). We attempted to match these variables in a relative sense using pre-placement calculations and ultrasound guidance, which improved upon previous work by quantifying true catheter tip depth; however, relative depth still varied between participants. Future studies could refine these methods to further reduce between-participant variance in placement. Third, the time synchronization between the Doppler ultrasound and ROM data was not sufficient to allow contraction phase- and ROM-specific blood flow analyses. We proposed that at some loads there may be higher blood flow during the eccentric compared to concentric muscle actions, but without direct phase-specific measurements, this remains speculative. Fourth, the non-failure exercise bouts involved different numbers of repetitions, which complicates interpretation of the post-exercise hyperaemia results. We cannot separate the potential effect of the additional repetitions performed at 20% and 30% 1RM on the post-exercise hyperaemic response. Repetitions were not matched between loads so that each load could reach its minimum SmO_2_ value, similar to previous work (8), as more repetitions are required at lower loads to do so. Finally, EMG was recorded from the VM and IMP within the VL, owing to limited space on the VL for simultaneous IMP, NIRS, and EMG measurement. While not ideal, we believe these results are still representative of the quadriceps, but future work should collect EMG and IMP from the same muscle, and, if possible, at approximately the same location.

## Conclusion

There are marked differences in blood flow responses below and above the COT based upon differences in contraction-induced IMP. Exercise against loads ≥COT elicits higher mean IMP values, leading to a reduction in exercising blood flow and muscle oxygenation, and a large increase in the post-exercise hyperaemic to exercising blood flow ratio compared with loads <COT. Together, these results support the COT as a relevant physiological concept defined by contraction-induced intramuscular microvascular occlusion. IMP was also found to have strong positive relationships with relative load, absolute load, and normalised EMG at both the group and individual levels. Finally, reconciling the above IMP relationships with load and normalised EMG, we propose that IMP, while related to load, is more likely a product of total motor unit recruitment and firing rate, given similar mean IMP values at task failure in the higher- and lower-load exercise bouts.

