## Supplementary Figure 1 for "Leg blood flow and muscle oxygenation are reduced when exercising with loads ≥40% 1RM compared to loads ≤30% 1RM due to increases in contraction induced intramuscular pressure"

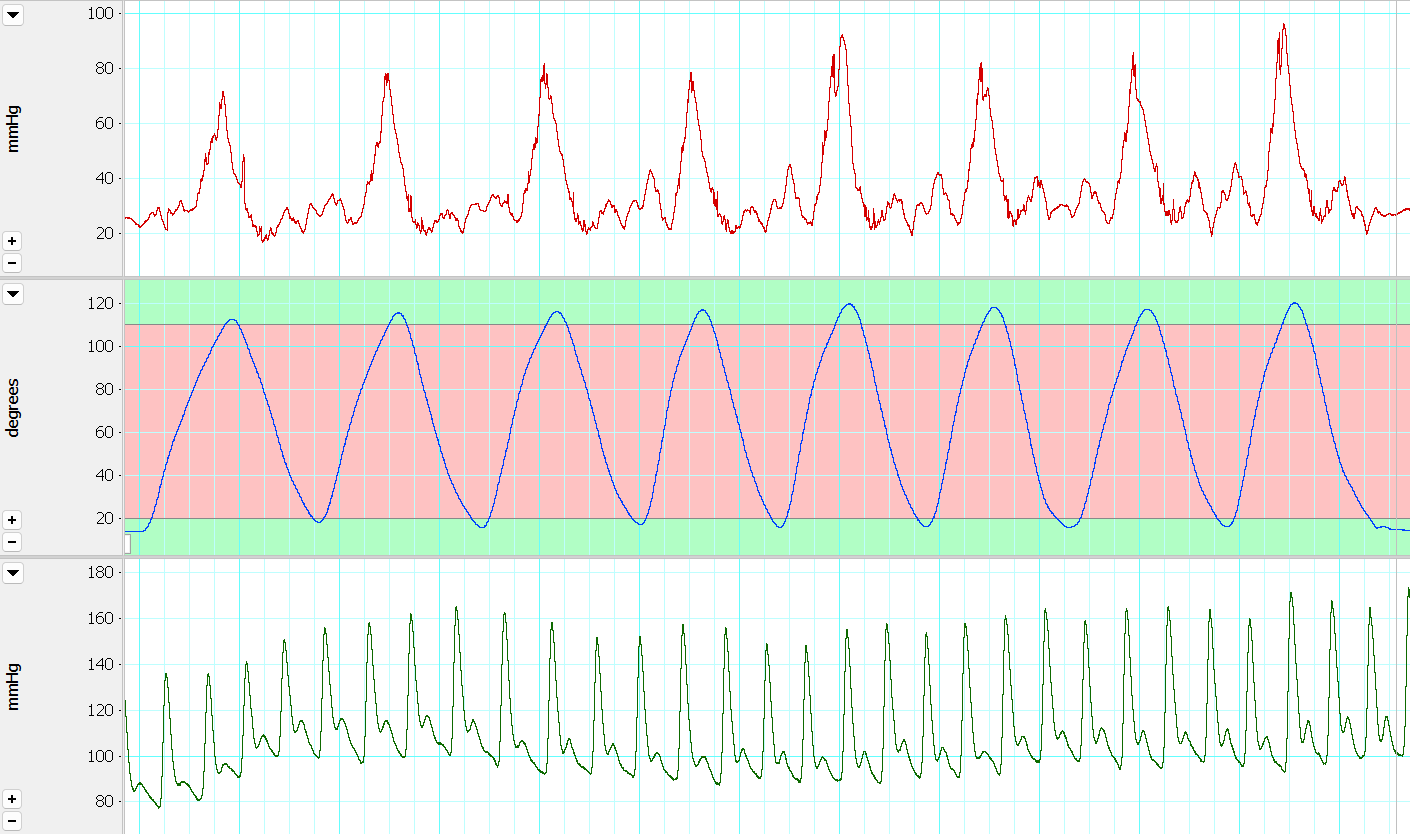

Supplementary Figure 1 – Representative LabChart data collection

This sample data is from one participant completing eight repetitions at 30% 1RM. The top trace is IMP (red line), the middle trace is joint angle with green zones representing the top and bottom ranges requires for a full range of motion (blue line), and the bottom trace is blood pressure (green line).
