## Supplementary Figure 2 for "Leg blood flow and muscle oxygenation are reduced when exercising with loads ≥40% 1RM compared to loads ≤30% 1RM due to increases in contraction induced intramuscular pressure"

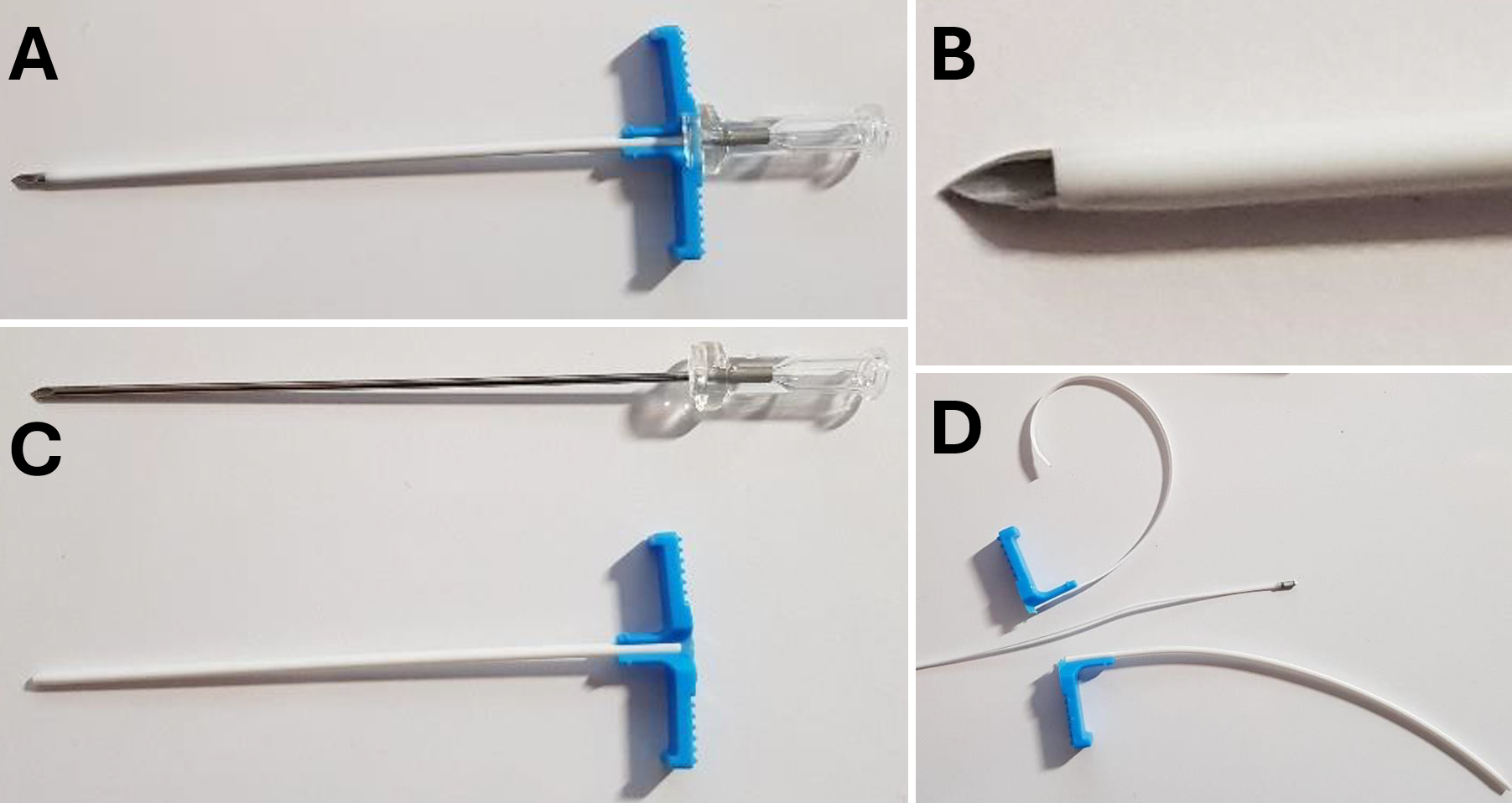

Supplementary Figure 2 - Introducer needle components. Visual of the introducer needle components (A, B, C) as well as how the cannula can be removed from around the catheter to allow for disposal.
