## Supplementary Figure 3 for "Leg blood flow and muscle oxygenation are reduced when exercising with loads ≥40% 1RM compared to loads ≤30% 1RM due to increases in contraction induced intramuscular pressure"

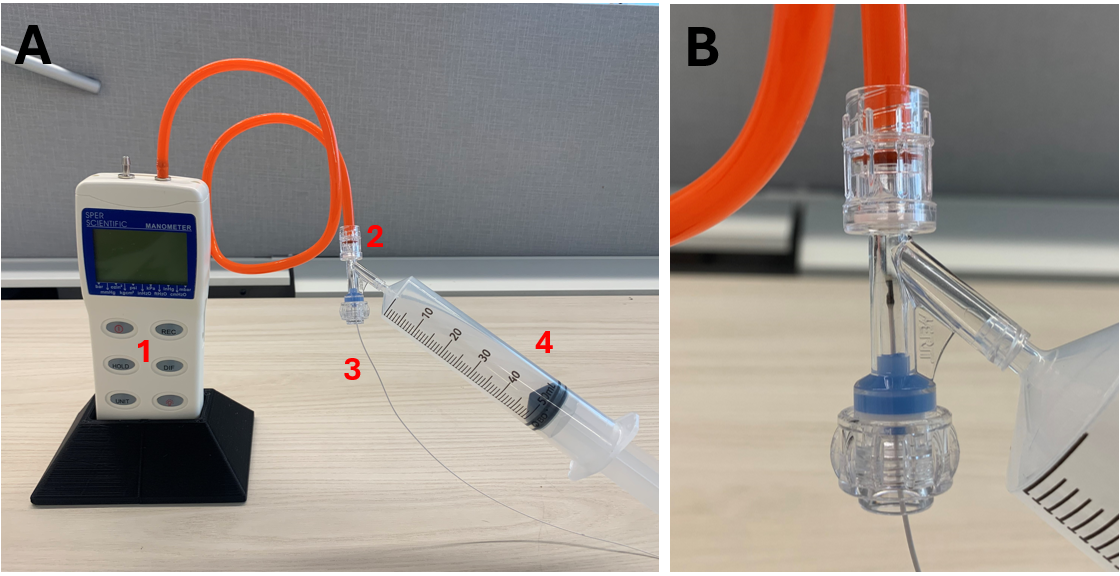

Supplementary Figure 3 - IMP catheter calibration setup. Visual setup of the IMP catheter calibration equipment. Panel A depicts the manometer (1), hemostasis y-valve connector (2), IMP catheter (3), and 50 mL syringe (4). Panel B provides a zoomed in image showing the connection of all components.

**IMP Catheter Calibration Methods**

IMP catheters were calibrated after removal from the participant, as calibrating before insertion would have compromised catheter sterility. Following removal, the catheters were reconnected to the recording setup so that the calibration values were captured in each participant’s LabChart file. Calibration required the use of a hemostasis Y-valve connector (V-01, 97380, Hemostasis Valve Y Connector, Rotating Male Luer lock, Female Luer lock Sideport, Qosina Corp., Ronkonkoma, NY, USA), which had a Tuohy-Borst valve, a Luer lock side port, and a distal connecting port (Supplementary Figure 4). The IMP catheter was inserted through the Tuohy-Borst valve and secured in place by twisting the valve shut. A manometer (Manometer 30 PSI, Sper Scientific Direct, Scottsdale, AZ, USA) was connected to the distal connecting port and set to display in mmHg. Calibration began by recording the pressure experienced by the catheter with no externally applied pressure; this value was taken 0 mmHg. Then, a 50 mL Luer lock syringe was attached to the side port of the Y-valve connector, and a member of the research team applied pressure to the catheter in 100 mmHg increments. Specifically, they syringe plunger until 100, 200, 300, and 400 mmHg were displayed on the manometer and held at each of these values for 5 s; the steadiness of each hold was monitored visually using the LabChart pressure trace. Once each pressure value had been recorded, the LabChart file was stopped, and a multipoint calibration was conducted using the Multipoint Calibration add-on from ADInstruments. Briefly, the manometer values were used to convert voltage to mmHg, and a linear fit was applied to each participant’s calibration, yielding an R^2^ of 0.98 or greater.
