## Supplementary Figure 4 for "Leg blood flow and muscle oxygenation are reduced when exercising with loads ≥40% 1RM compared to loads ≤30% 1RM due to increases in contraction induced intramuscular pressure"

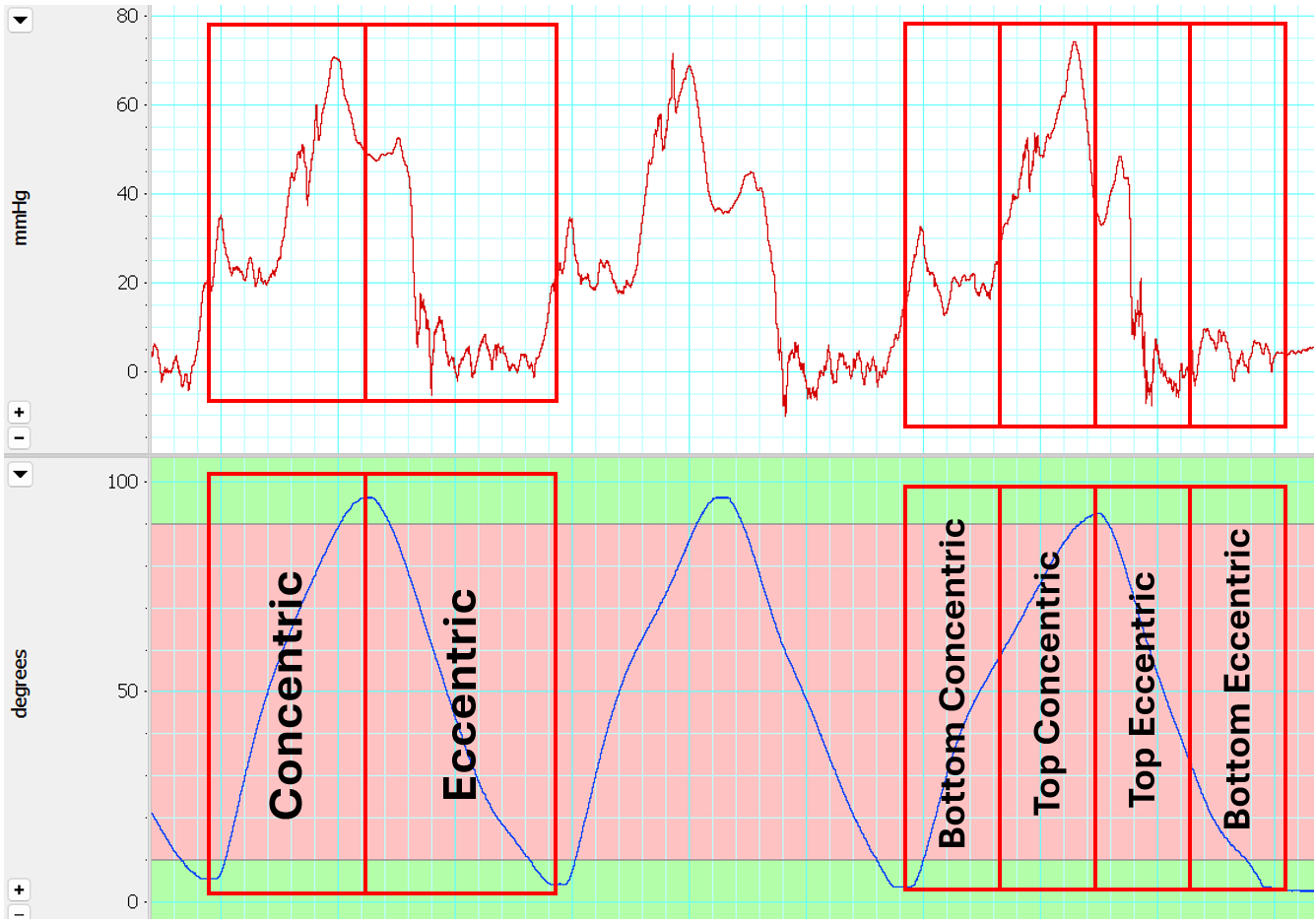

Supplementary Figure 4 - Representative IMP and kicking arm position data. Depicted are three sample repetitions. The top trace represents IMP, and the bottom trace represents joint angle. Labels have been placed on the joint angle trace to showcase the concentric and eccentric actions, as well as how the repetitions were split into bottom concentric, top concentric, top eccentric, and bottom eccentric for analysis.
