## Supplementary Figure 5 for "Leg blood flow and muscle oxygenation are reduced when exercising with loads ≥40% 1RM compared to loads ≤30% 1RM due to increases in contraction induced intramuscular pressure"

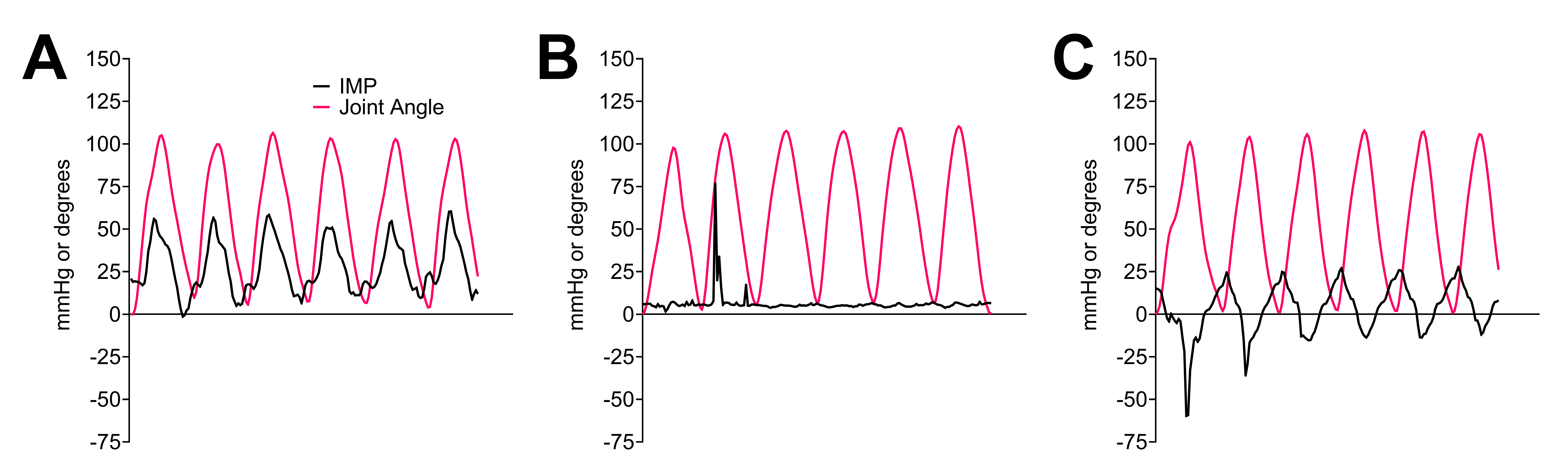

Supplementary Figure 5 - Representative data of expected and non-physiological IMP responses. Expected IMP and joint angle relationship (A) compared to a non-physiological IMP response (B) and an inverted IMP response (C).
