## Supplementary Figure 6 for "Leg blood flow and muscle oxygenation are reduced when exercising with loads ≥40% 1RM compared to loads ≤30% 1RM due to increases in contraction induced intramuscular pressure"

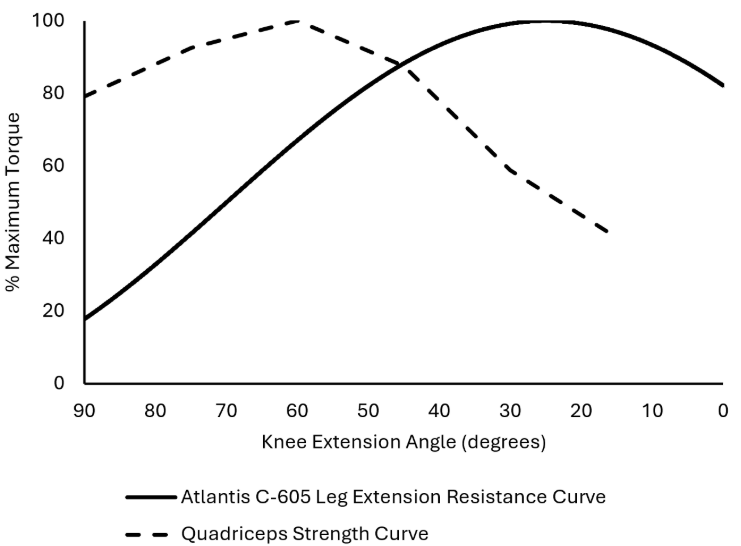

**Supplementary Figure 6 -** Comparison of the Atlantis C-605 Leg Extension resistance curve and quadriceps strength curve from Kulig et al. 1984 (39) across a 90°-0° range of motion. The resistance curve for the C-605 leg extension was calculated using trigonometry.
